# Porphyrin-driven redox tuning in structurally defined *de novo* heme proteins

**DOI:** 10.64898/2026.06.09.731085

**Authors:** Cameron Mellor, Christopher Williams, Ethan L. Bungay, Jessica C. Berrones-Reyes, Rob Barringer, Catherine R. Back, Paul Molinaro, Ronald L. Koder, Bruce R. Lichtenstein, Adrian J. Mulholland, Matthew P. Crump, J. L. Ross Anderson

## Abstract

Designing redox proteins with predictable and tuneable electron transfer properties is a major goal in *de novo* bioenergetics. Here we show that replacing heme B with a series of structurally conservative non-natural metalloporphyrins enables broad modulation of redox potentials over 400 mV in the *de novo* designed monoheme m4D2 and diheme 4D2 T19D. The non-natural porphyrins bind with high affinity and do not compromise either the heme binding site or global protein structure, as evidenced by X-ray crystallography and NMR spectroscopy. We also report the native-like NMR structure of m4D2 loaded with the non-natural and symmetric iron 2,4-dimethyldeuteroporphyrin IX, confirming our modular approach to tetrahelical redox protein design. This work establishes a versatile platform for constructing tuneable electron carriers for engineered bioenergetic pathways and bioelectronic applications.

## Introduction

Redox proteins are central to life, mediating electron flow through electron conducting respiratory, photosynthetic and metabolic assemblies and pathways, playing key roles in the generation and usage of cellular energy^1, 2^. Within these proteins and multiprotein assemblies, electrons move stepwise between redox active cofactors by quantum tunnelling, a process facilitated by the proteinaceous medium housing the cofactors^2, 3, 4^. The rates of these intercofactor hopping steps are determined principally by the distance between the cofactors, the reorganisation energy (associated with reorientation of dipoles local to redox cofactors), and the driving force for electron transfer (given by the relative redox potentials of the donor and acceptor). Additionally, at short intercofactor distances (<6 Å), orientation-dependent electronic coupling becomes significant, especially for conjugated cofactors such as heme^5^.

If the intention is to build *de novo* electron-conducting proteins and assemblies, it is necessary for a designer to understand and control these parameters within tractable^6, 7^ and structurally well-defined proteins^8, 9^. To simplify the *de novo* design process and focus on functional sufficiency at the expense of optimisation, it may be desirable to exclusively focus on intercofactor separation and the free energy of electron transfer within such *de novo* proteins^4^; for single cofactor-containing proteins, this can be further reduced to controlling redox potential at relatively accessible cofactor binding sites. Ultimately, such simple *de novo* proteins could serve as tuneable, soluble and diffusible electron carriers for interaction with other redox proteins and enzymes, whether natural or designed.

Excluding chromophores, cofactors and reactive cofactor intermediates that form high energy photoexcited or radical states, nature typically accesses a window of redox potential in proteins between -700 and +800 mV^10, 11^ through redox cofactors such as hemes, iron-sulfur clusters, flavins and metal ions. Considering hemes alone, this range reduces to -550^12^ to +385^13^ mV in the reported literature, though this still represents 935 mV of accessible redox potential available for redox engineering. While delineating the exact determinants of redox potential for a given protein remains challenging, it is understood that axial ligation, secondary coordination sphere, solvent accessibility, local electrostatic environment and covalent heme modifications play significant roles^10, 14^. Indeed, we, and others, have demonstrated that minor changes to these features, such as manipulating axial ligand hydrogen bonding networks, mutating axial ligands and altering electrostatic environments^9, 15, 16, 17-19^, can result in modulation of heme redox potentials that can often be predicted with some accuracy^9, 15, 18^.

*De novo* protein design has proved valuable in probing and understanding these effects, and there are many reports where bioenergetic proteins of reduced complexity, typically called maquettes, have been successfully designed and characterised^6, 8, 20, 21^, albeit most exhibiting poorly defined three-dimensional structures^20, 22^. Recently, we have described the modular design of mono-, di- and tetraheme maquettes (named m4D2, 4D2 and e4D2 respectively) with significantly improved structural resolution over previous mono- and multiheme maquettes^9, 18^. These proteins are built on the four α-helix bundle, a common natural backbone that provides a versatile and robust chassis with excellent capacity for design both in soluble^6, 8, 9, 18, 20^ and transmembrane^23, 24^ environments. Though we were able to demonstrate some redox control over the monoheme m4D2^9, 18^, we wished to further expand the redox range within m4D2 and our other related multiheme proteins. This would enable us to then select variants with redox potentials matching potential interaction partners or electron transfer pathways downstream, while ideally retaining the well-folded, native-like structure of the original protein^9^.

Here we describe the versatility of the heme binding sites in the monoheme m4D2 and the T19D mutant of the diheme 4D2 in accommodating non-natural porphyrin (NNP) analogues of heme B with redox-potential altering substitutions in the 2- and 4-positions of the conjugated porphyrin ring (**Figure 1**). Using these NNPs, we were able to access 400 mV of redox potential without redesigning or significantly affecting the overall protein fold; crystal structures of 4D2 T19D loaded with two of the NNPs show minimal global or local structural perturbation, and 2D NMR spectra of all tested m4D2-NNP variants feature excellent peak dispersion patterns that are consistent with our previously reported heme B-bound spectrum^9^. The NNPs all bind to the two *de novo* proteins with nanomolar affinity and exhibit varying binding kinetics that appear complex. We also report a solution NMR structure of m4D2 with a bound symmetric NNP, confirming the original design and highlighting the restricted dynamics of the heme-bound protein. Taken together, these results highlight the robust nature of these *de novo* proteins and indicate their potential in acting as readily tuneable electron carriers for incorporation into natural or artificial electron transfer pathways.

**Figure 1.**
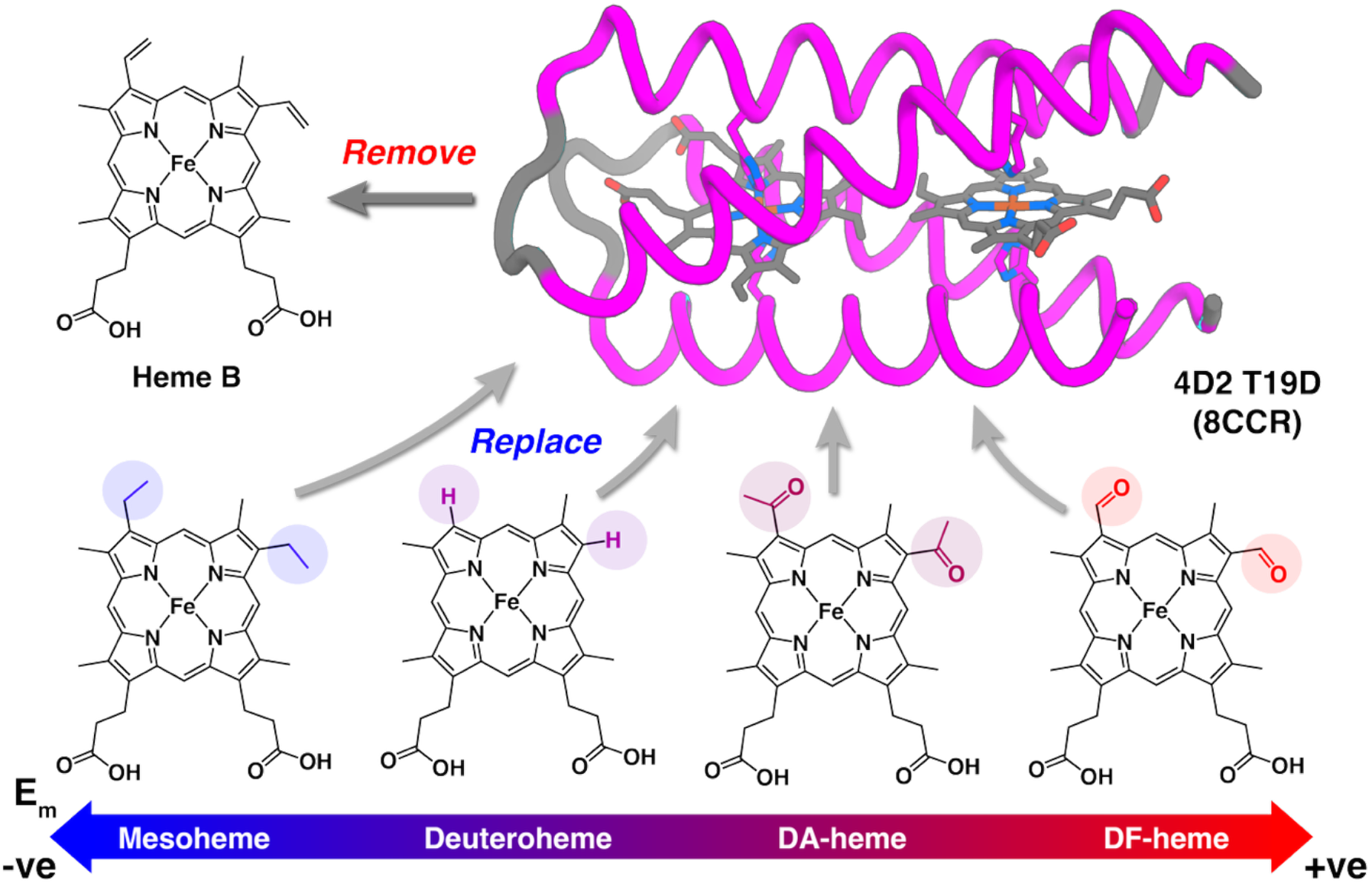
Facile heme replacement in m4D2 and 4D2 T19D with structurally conservative NNPs facilitates broad redox tuning and selection. Heme B is removed in these proteins through acid:butanone extraction, and can be replaced by NNPs with structurally modest, yet electronically affecting, substitutions in the 2- and 4-positions.

## Results and Discussion

To probe the ability of our *de novo* protein heme binding sites to accept NNPs, we selected two designs with well-defined structures for investigation: the T19D mutant of 4D2 which proved to be readily crystallisable with heme B bound; and m4D2, for which we previously reported 2D NMR spectra with excellent peak dispersion when loaded with the symmetric, non-natural iron 2,4-dimethyldeuteroporphyrin IX (DM-heme)^9^. It should be noted that the heme binding site of m4D2 is identical in sequence to the equivalent site in 4D2 T19D. For the porphyrins, we selected NNPs with substitutions in the 2 and 4 positions of the tetrapyrrole ring (**Figure 1**), thereby retaining key protein:ligand binding interactions such as the heme propionate:arginine ion pairs. Previous work by Lu^17^, and others^25, 26^, has revealed the ability of natural and engineered proteins to accept such porphyrins – so long as the porphyrin substitutions are relatively conservative – with minimal structural perturbations to the protein, while unlocking a broad range of redox potential^17, 21, 27^. For this study, we selected NNPs with either protons (iron deuteroporphyrin IX; deuteroheme), ethyl (iron mesoporphyrin IX; mesoheme), acetyl (iron 2,4-diacetyldeuteroporphyrin IX; DA-heme) or formyl (iron 2,4-diformyldeuteroporphyrin IX; DF-heme) groups in the 2,4 positions, thus providing direct electronic modulation of the porphyrin ring through each group’s differing electronic properties.

We initially examined the binding of these selected NNPs to our apoproteins through simple binding titrations monitored by electronic spectroscopy and fit to the quadratic tight binding equation (**Figure 2, Table S1**). All NNPs bound stoichiometrically to both proteins with nanomolar affinities, albeit with generally lower affinity than heme B^9^ (**Figure 2, Table S2**). The weakest binding to m4D2 is exhibited by DF-heme and DA-heme (*K*_*D*_ = 15.6 nM & 12 nM), presumably as a consequence of adding both steric bulk into the binding site (acetyl methyl group of DA-heme) and the carbonyls acting as unsatisfied hydrogen bonding acceptors in the hydrophobic core of the protein. Except for DF-heme, it is also apparent that there is a cumulative, destabilising effect when more than one NNP is bound to the protein; the binding affinity decreases by approximately 2-fold or more for DA-heme, mesoheme and deuteroheme in 4D2 T19D versus m4D2, though there is no evidence of negative cooperativity in any of the binding titrations. However, it is well understood that the errors intrinsic to this technique and fitting equation can be significant^28^, and while the data indicate stoichiometric and high affinity (*K*_*D*_ <40 nM) binding, the granularity we observe may not accurately represent the true variance in dissociation constants across the NNPs.

**Figure 2.**
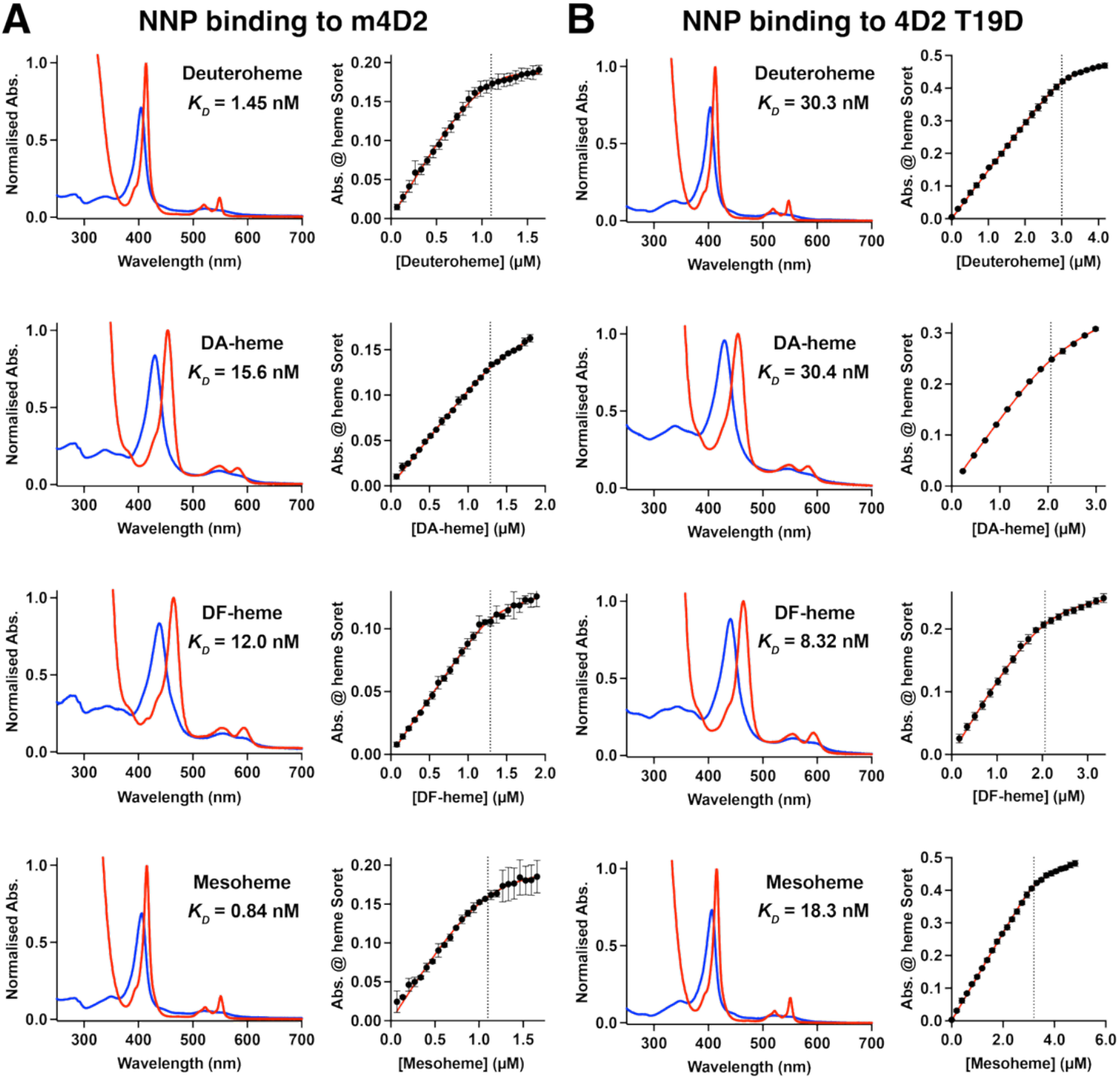
Spectroscopic and binding analyses of NNPs to *de novo* heme proteins. (A) Ferric (blue) and ferrous (red) UV/visible spectra of apo-m4D2 with NNPs, and binding isotherms of apo-m4D2 with ferric NNPs. (B) Ferric (blue) and ferrous (red) UV/visible spectra of apo-4D2 T19D with NNPs, and binding isotherms of apo-4D2 T19D with ferric NNPs. UV/visible spectra are normalised to the ferrous Soret (γ) absorption band and recorded from 5-10 μM NNP:protein in 20 mM CHES, 100 mM KCl, pH 8.6. Binding isotherms were recorded at 1-3 μM apoprotein in 20 mM CHES, 100 mM KCl, pH 8.6. Error bars represent standard deviations for each data point in the binding isotherms, and the dashed lines indicate the point where [NNP] = [protein].

To gain further understanding of the binding process in our simplest protein chassis, we subsequently probed the kinetics of NNP binding to m4D2 using stopped flow spectrophotometry, mixing stoichiometric quantities of the ferric NNPs with apo-m4D2 (**Figure 3**). Under these conditions, the binding kinetics are surprisingly complex, especially for heme B which exhibits pronounced, multiphasic binding kinetics. In fact, only the DF-heme and DA-heme kinetics can be described using a biphasic model, with mesoheme and deuteroheme also exhibiting a kinetic complexity that confounds simple interpretation. We suggest these observations may indicate a multistep mechanism of porphyrin de-aggregation, entry to the protein core and bis-histidine ligation within a conformationally heterogeneous binding site, as previously suggested by Solomon *et al*^29^. Evidence for the latter is provided by the 2D ^1^H-^15^N HSQC NMR spectrum of apo-m4D2^9^, which, while demonstrating relatively good peak dispersion, also indicates the presence of some conformational heterogeneity. This is consistent with the design of m4D2^9^, and PS1 from the Degrado group^30^, in which a rigid, well-packed protein module was placed adjacent to a more flexible metalloporphyrin binding module that could accommodate the ligand with a low kinetic barrier to entry. Qualitatively, in m4D2, mesoheme binds most rapidly, followed in order by deuteroheme, heme B, DA-heme and DF-heme, the latter requiring approximately 8 minutes to occupy all vacant binding sites. In contrast, mesoheme had completely occupied the available binding sites within 2 seconds. While a relationship between NNP hydrophobicity and binding kinetics to a *de novo* protein has been proposed^29^, it is hard to disentangle the various contributions of the 2,4 substitutions studied here on the other possible mechanistic steps, and no simple correlation between NNP partition coefficients and the binding kinetics is apparent.

**Figure 3.**
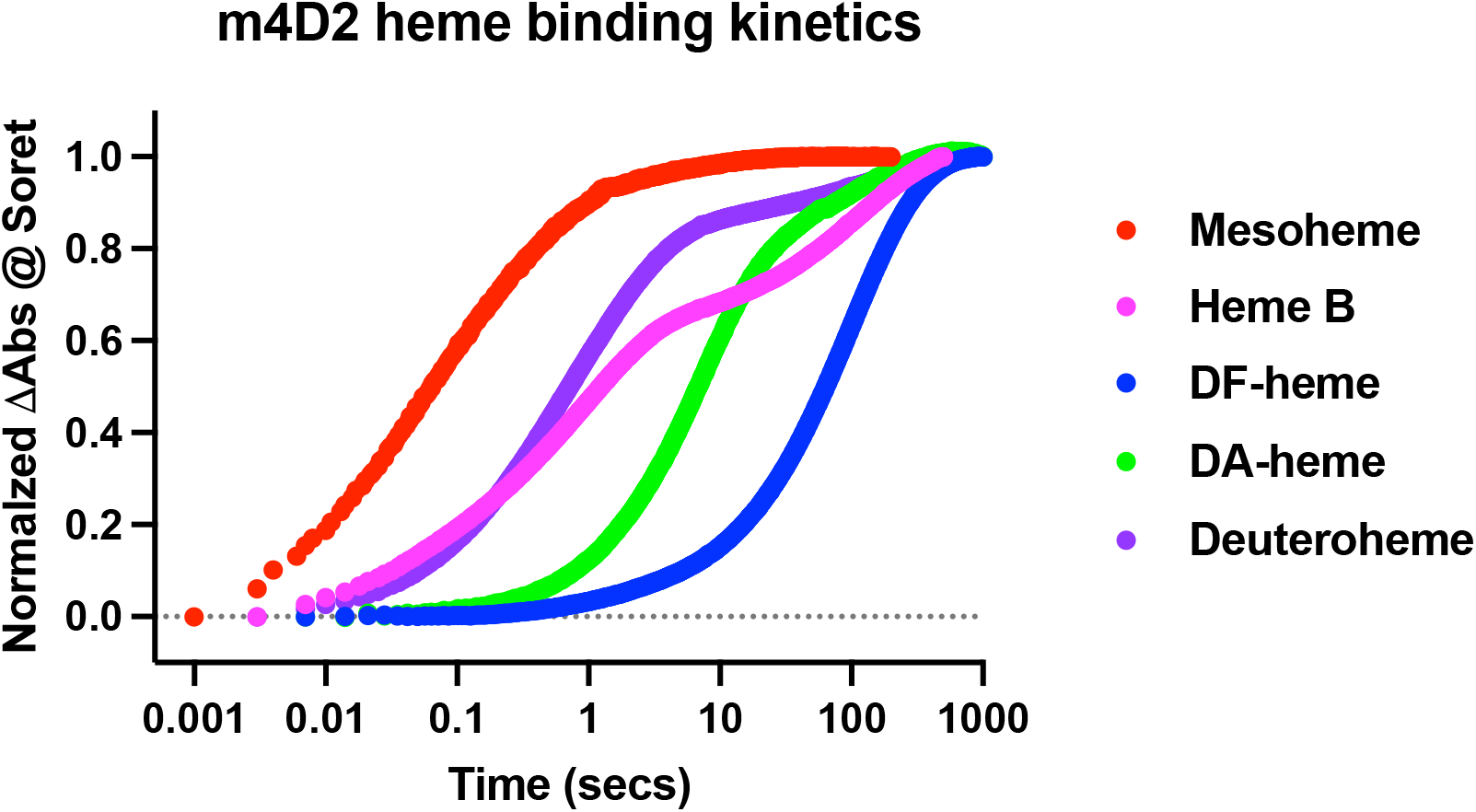
NNP:m4D2 binding kinetics assessed by stopped-flow spectrophotometry. Normalised kinetic traces representing the absorbance changes at the ferric NNP Soret band primarily exhibit complex multiphasic kinetics while demonstrating significant differences in overall binding rates. Data was recorded rapidly mixing equimolar concentrations of apo-m4D2 and NNP (both 10 μM) in 20 mM CHES, 100 mM KCl, pH 8.6.

With NNP binding established, we used circular dichroism spectroscopy to probe secondary structure and thermal stability of the m4D2:NNP and 4D2 T19D:NNP complexes (**Figure 4**). All m4D2:NNP variants exhibited identical CD spectra and thermal scan data indicative of highly thermostable proteins that lose only a small degree of helicity up to 95°C. These data are also identical to those that we previously reported for the m4D2:heme B complex^9^, indicating minimal structural perturbations on substituting heme B for the NNPs, as well as a stabilising effect on the apoprotein when the porphyrin binds. The T19D mutant of 4D2 has a clear destabilising effect on the apoprotein relative to the original 4D2 design, with no discernible secondary structure from 20°C to 95°C (**Figure S1**). In contrast, apo-4D2, while possessing a low melting transition (*T*_*m*_) at approximately 25°C, appeared predominantly helical at low temperature^9^. It is therefore likely that placing the negatively charged aspartate side chain in the core of T19D 4D2 without a paired positively charged residue is responsible for this destabilisation. With heme B bound, T19D exhibits similar thermostability to holo-4D2 and NNP-bound m4D2 (**Figure S1**); however, with NNPs bound, there is a notable loss in thermal stability, with *T*_*m*_ values between 70 and 86°C (**Figure 4**). Consistent with the binding titrations, DA-heme and DF-heme complexes with T19D exhibit the lowest *T*_*m*_ values (72 and 70°C respectively), highlighting the destabilising effect of additional steric bulk and unsatisfied hydrogen bonding in the hydrophobic core on global protein stability.

**Figure 4.**
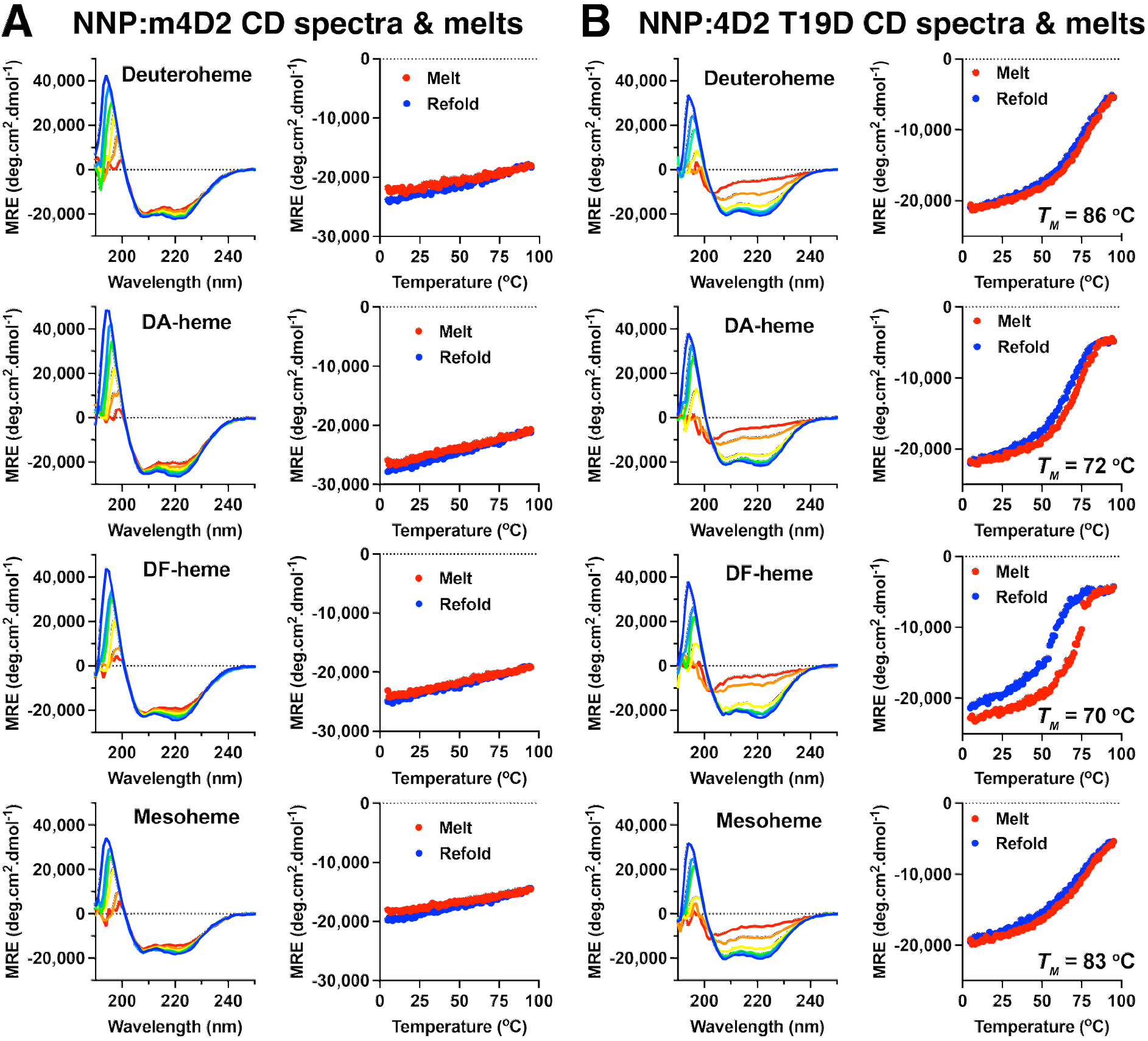
Circular dichroism spectra and thermal melts of NNP:protein complexes. (A) Variable temperature far-UV circular dichroism spectra and temperature-dependent CD signal at 222 nm for m4D2:NNP complexes. (B) Variable temperature far-UV circular dichroism spectra and temperature-dependent CD signal at 222 nm for 4D2 T19D:NNP complexes. CD spectra were collected on 1-3 μM NNP:protein in 20 mM CHES, 100 mM KCl, pH 8.6, and full spectra at 5, 25, 35, 55, 75 and 95 °C are coloured blue, light blue, green, yellow, orange and red respectively. For the temperature dependent 222 nm CD signal traces, red represents the thermal melt from 5 to 95 °C, while blue represents the corresponding cooling of the protein from 95 to 5 °C. Melting temperatures (*T*_*M*_) are indicated for 4D2 T19D:NNP complexes.

To further probe the effects of NNP binding to these *de novo* proteins, we attempted to crystallise all NNP-bound versions of 4D2 T19D and were successful in obtaining crystals for 4D2 T19D:mesoheme (PDB 9H4C) and 4D2 T19D:DF-heme (PDB 9SY9) (**Table S3**). Using molecular replacement to solve the X-ray diffraction datasets collected at Diamond Light Source, we successfully solved the crystal structure of the mesoheme variant at 1.98 Å resolution, and the DF-heme variant at 2.0 Å resolution (**Figure 5**).

**Figure 5.**
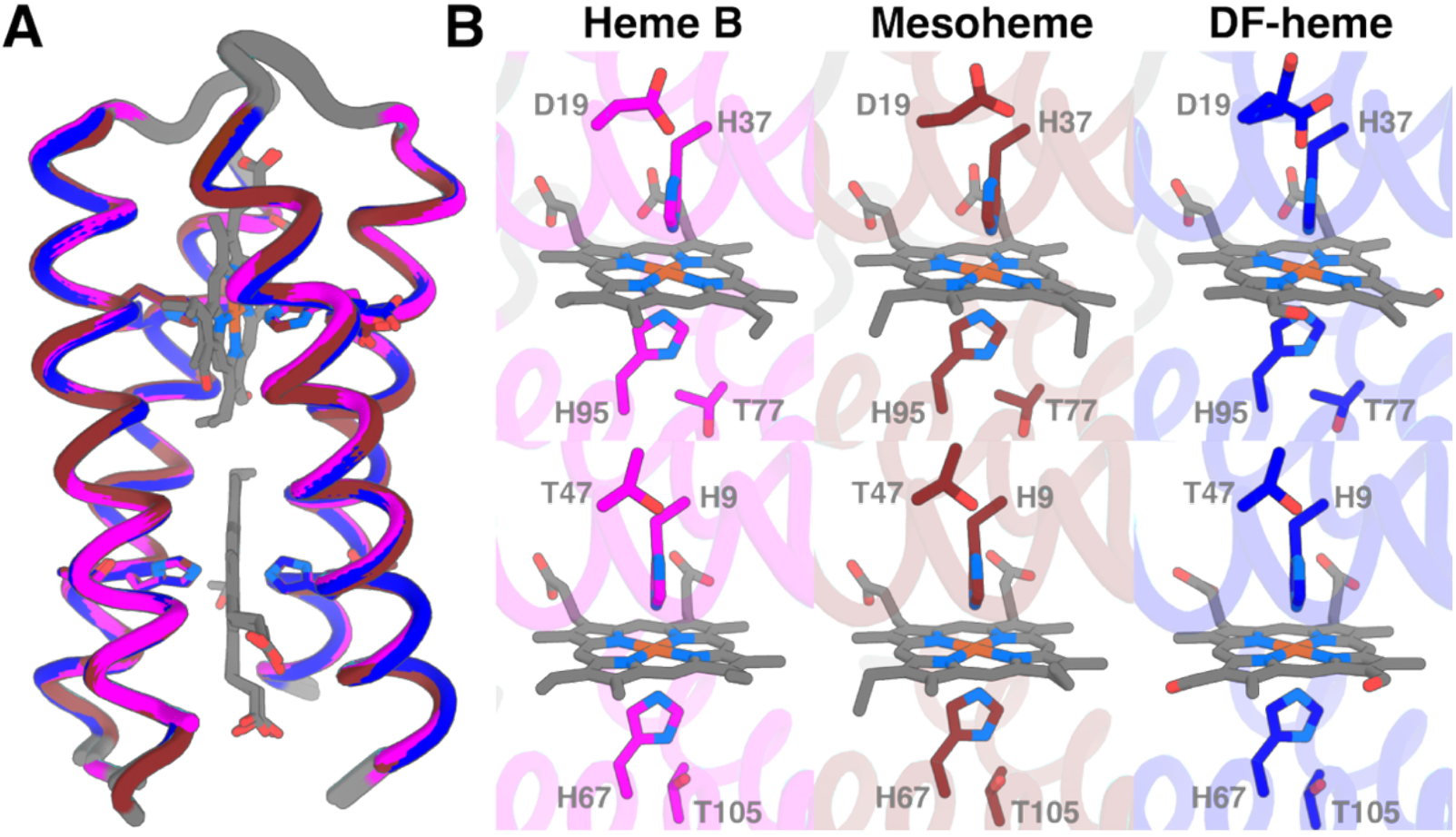
Replacing heme with NNPs in 4D2 T19D causes minimal structural perturbations to the protein. (A) Overlay of the crystal structures for 4D2 T19D with bound heme (PDB 8CCR; magenta), mesoheme (PDB 9H4C; chocolate) and DF-heme (PDB 9SY9; blue). (B) Views of the heme binding sites within 4D2 T19D loaded with heme B, mesoheme and DF-heme.

These new structures align exceptionally well to the original 4D2 T19D:heme B structure^9^, exhibiting all-atom RMSD values of 0.260 Å and 0.248 Å for mesoheme and DF-heme variants respectively, with no major backbone, sidechain or porphyrin conformational differences, or crystallographic B-factors (**Figure S2**) evident. This is perhaps unsurprising given that vinyl, formyl and ethyl groups are essentially isosteric, though the ethyl groups should exhibit greater conformational flexibility with free rotation around the saturated carbon-carbon bonds. While we were unable to crystallise the other 4D2 T19D:NNP complexes, we obtained 2D ^1^H-^15^N HSQC spectra for each m4D2:NNP complex (**Figure S3**). All four spectra exhibit good signal dispersion consistent with, and nearly identical to, the analogous spectrum of heme B bound to m4D2^9^, with the DF-heme spectrum exhibiting fewer peaks and better overall resolution than the others. We highlight that in the m4D2:NNP spectra, there are more signals than would be expected for a protein of this size, which we have previously ascribed to the asymmetry of the heme and a lack of orientational selection from the binding site itself^9^. This is particularly apparent in our crystal structures of the diheme 4D2 and 4D2 T19D with heme B, which exhibit similar electron density in positions 1, 2, 3 and 4 of the tetrapyrrole ring and can be ascribed to a 180° rotation of the asymmetric heme B within the site. The structure of 4D2 T19D:mesoheme displays analogous behaviour, with weakly defined electron density for the mesoheme methyl and ethyl groups at the double loop heme binding site, apparent in the 2F_0_-F_c_ OMIT map (**Figure S4**). Stronger density for these groups can be observed at the termini heme site, indicating some preference for one conformation for mesoheme within this protein. Conversely, the equivalent 4D2 T19D:DF-heme structure reveals increased preference for a single porphyrin conformation in both heme binding positions, in agreement with the fewer and more resolved peaks of the NMR spectrum (**Figure S3**). We subsequently examined the electrostatic surface potential of the heme binding sites within this structure (**Figure S5**). The DF-heme at the termini site is positioned with its 4-formyl carbonyl group oriented towards a region of positive charge and decreased hydrophobicity due to the sidechain orientations of R64, R68 and Q49. The favourable interaction between the partial negative charge on the carbonyl oxygen and the positive electrostatic surface of the binding site, and the decreased hydrophobicity may therefore contribute to the orientational preference in this site, though it is less clear what would determine equivalent preference in the double loop site.

We previously reported that the 2D ^1^H-^15^N HSQC NMR spectrum of m4D2:heme B can be significantly improved by replacing heme B with a symmetric heme analogue, iron (III) 2,4-dimethyldeuteroporphyrin IX (DM-heme), resulting in a well-resolved spectrum^9^. To enable additional data collection and complete the structural assignment, we developed a new synthetic route to DM-heme from hemin chloride (**Scheme S1**). Following simultaneous demetallation and esterification to the protoporphyrin IX dimethyl ester, we oxidised the vinyl groups to the corresponding aldehydes in a two-step process using a modified version of the synthesis reported by Kahl *et al*^31^. We then used the Caglioti reaction to convert the aldehydes to methyl groups *via* tosylhydrazone adducts that we reduced under mild conditions^32^. We finally metalated the resulting porphyrin with iron chloride and hydrolysed the methyl esters^33^, obtaining the iron 2,4-dimethyldeuteroporphyrin IX at 12% yield after 7 steps.

Using this DM-heme, standard 3D triple-resonance and ^15^N-/^13^C-edited NOESY experiments were recorded to assign the protein backbone and side chains, and to determine the structure of m4D2:DM-heme (PDB 9I38). The presence of the paramagnetic iron centre caused significant signal broadening and bleaching of resonances proximal to the heme, resulted in a limited and largely ambiguous set of heme-protein NOEs. These restraints alone were insufficient to define the heme orientation during structure refinement, leading to residual rotation of the heme about the metal-histidine coordination axis within the binding pocket. The heme-binding site of m4D2 is identical in sequence to the corresponding site in the 4D2 T19D variant, and the bound isosteric hemes in the 4D2 crystal structures adopt near identical orientations within the binding pocket (**Figure 5**). Therefore, to restrain the DM-heme orientation in the NMR structure to conformations consistent with those observed in the homologous heme B structure, a small number of supplementary restraints derived from the 4D2 heme B crystal structure (PDB 7AH0) was incorporated into the calculation during the final stages of refinement. The resulting structural ensemble closely matches the overall topology and backbone of 4D2 (**Figure 6**)^9^, with an average all-atom RMSD of 1.436 Å across all m4D2 ensemble structures and a range between 0.967 and 2.197 Å. Perhaps unsurprisingly, the main structural differences between ensemble structures are in the loop and termini conformations, with only modest variations in backbone and sidechain positions in these regions. The redesigned hydrophobic core is well packed with aromatics (Trp & Phe) occupying the former heme binding site in orientations very close to those specified in our original computational design model^9^ and in an AlphaFold3-derived structure prediction^34^ (**Figure S6**).

**Figure 6.**
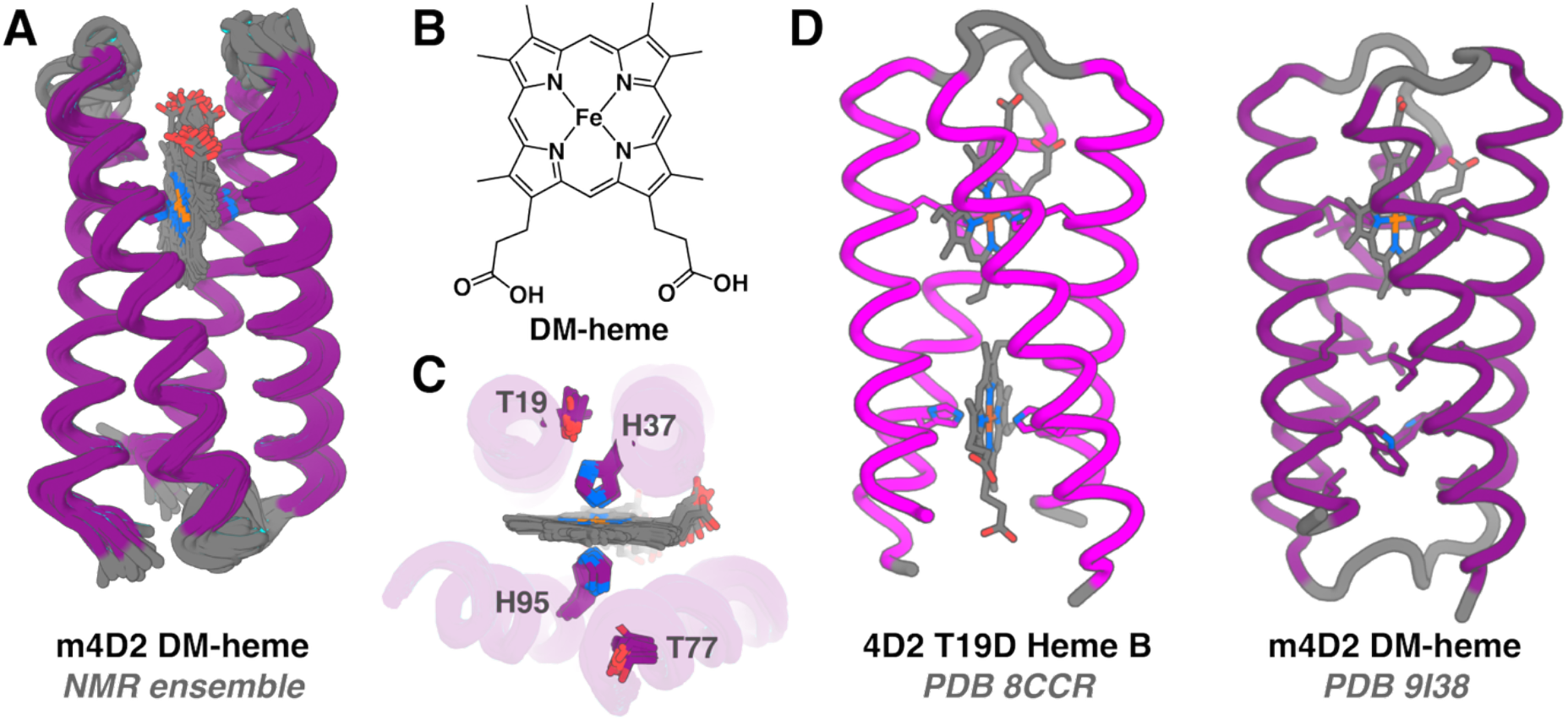
Structural analysis of the m4D2:DM-heme complex by NMR spectroscopy. (A) Ensemble (9I38) representing the 20 lowest energy solutions to the NMR spectral analysis of the m4D2:DM-heme complex. (B) Chemical structure of DM-heme. (C) DM-heme ligation and secondary H-bonding between binding site threonines and histidines (T19, 77 and H37, 95) across the ensemble. (D) Comparison of the diheme B 4D2 T19D (PDB 8CCR) crystal structure with a low energy structure from the m4D2:DM-heme complex NMR ensemble (PDB 9I38).

Following this structural characterisation and verification, we used optically transparent thin layer electrochemistry^35^ to measure redox potentials of the m4D2 and 4D2 T19D NNP complexes (**Figure 7**). For the single site m4D2, substitution of heme B with the series of NNPs results in a redox potential range of 376 mV between the mesoheme (-235 ± 1 mV) and DF-heme (+141 ± 7 mV). These redox potential shifts are consistent with previous work examining NPP substitution in natural and engineered proteins^17^. Given the distribution of potentials of the 5 NNPs tested, it is possible to select the potential of the protein within approximately 100 mV intervals, with further fine tuning available either through protein modification as previously reported^9, 18, 19^, or through further porphyrin modification. For the latter, this would require similarly modest synthetic modifications to the NNPs to avoid any deleterious effects on binding affinity. A possible route to achieve this, or to further expand the redox potential range, could be the selective and cumulative substitution of porphyrin protons with electron-withdrawing fluorines, either directly at the *meso* methines or on substituents situated at the 1-4 *β* positions.

**Figure 7.**
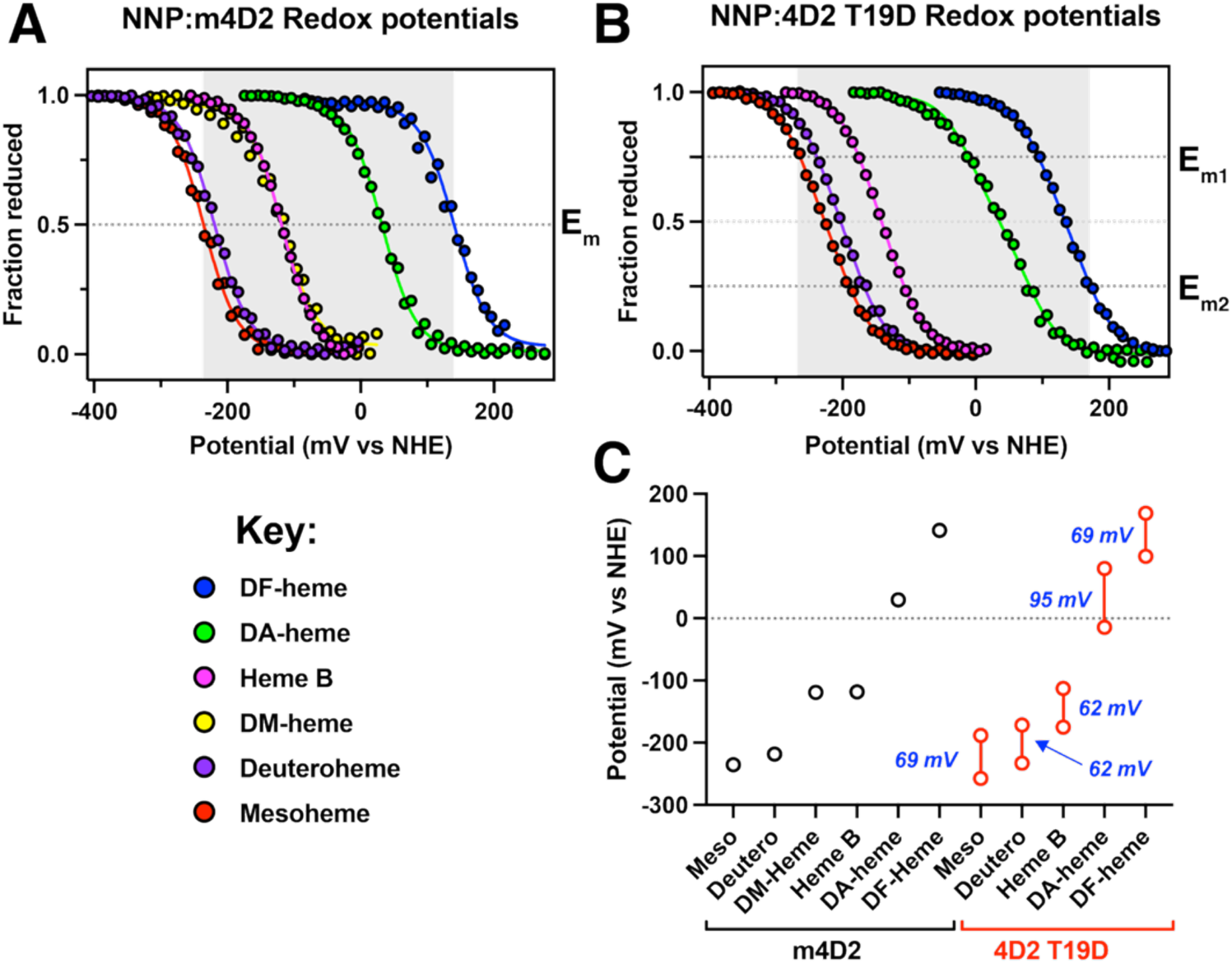
Redox analysis of the protein:NNP complexes reveals a broad and readily accessible window of redox potential. (A) Redox potentiometry of m4D2:NNP complexes recorded in 20 mM CHES, 100 mM KCl, pH 8.6. Data was recorded in triplicate and fitted to a single electron Nernst model. The Key provided below identifies the NNPs associated to the data. (B) Redox potentiometry of 4D2 T19D:NNP complexes recorded in 20 mM CHES, 100 mM KCl, pH 8.6. Data was recorded in triplicate and fitted to a sequential two electron Nernst model. (C) Fitted potentials for both m4D2:NNP and 4D2 T19D:NNP reveal over 400 mV of redox tuning available to the designer. Redox cooperativities in the 4D2 T19D:NNP complexes are indicated in blue on the figure.

Within the two-heme binding 4D2 T19D, the redox potentials of the 4 tested NNP complexes span 426 mV, from -257 ± 4 mV (mesoheme) to +169 ± 2 mV (DF-heme), which is greater than the equivalent m4D2 set. This further expansion is due, in part, to the redox cooperativity arising from both direct heme-heme electrostatic coupling and environmental electrostatic changes arising when the heme with intrinsically higher potential is reduced^18^. For this series of NNP complexes, this coupling is generally consistent in the range of 60-70 mV, with DA-heme 4D2 T19D exhibiting a slightly higher value of 95 mV, closer to that observed in the equivalent transmembrane version of 4D2^18, 23^. The origin of this increase relative to the other NNP complexes is not immediately clear and will require the acquisition of a high-resolution structure to unequivocally determine whether the acetyl groups impose structural perturbations not observed in the other NNP:protein complexes.

## Conclusions

These results further showcase the utility of our *de novo* protein scaffolds to act as versatile, tuneable bioenergetic components. We have demonstrated that at least 426 mV of redox potential is readily accessible within an unaltered heme binding site with minimal structural perturbation and the maintenance of a native-like state. While similar heme substitutions have been successfully performed on natural^25^, engineered^17, 26^ and *de novo* heme-containing maquettes^21, 27^, the latter group has typically suffered from structural heterogeneity that precluded the acquisition of high resolution structural data, therefore hindering any downstream atomistical redesign. Here, the option to redesign remains available, and future efforts will focus on engineering protein:protein interactions with *de novo* and/or natural electron transferring proteins to facilitate rapid inter-molecular transport in artificial or semi-artificial redox circuitry.

## ASSOCIATED CONTENT

### Supporting Information

The following files are available free of charge:

- Materials and methods; synthetic scheme for Fe(III) 2,4-dimethyl-deuteroporphyrin IX; UV/visible spectroscopic data for m4D2:NNP and 4D2 T19D:NNP complexes; NNP binding affinities to m4D2 and 4D2 T19D; crystallographic data collection and refinement statistics for 4D2 T19D:mesoheme and 4D2 T19D:DF-heme structures; redox potentials of m4D2:NNP and 4D2 T19D:NNP complexes measured by redox potentiometry; biophysical analysis of m4D2 T19D:heme B; crystallographic analysis of 4D2 T19D:NNP/heme variants; NMR analysis of NNP binding to m4D2; 2F_O_-F_C_ OMIT maps of NNP complexes with 4D2 T19D; figure rationalising orientational preference of DF-heme in 4D2 T19D; NMR structural ensemble of m4D2:DM-heme and comparison with computational structural predictions; HPLC chromatograms of 2,4-Diformyl-deuteroporphyrin IX dimethyl ester *p*-ditosylhydrazone.

## AUTHOR INFORMATION

### Authors

**Cameron Mellor** – School of Biochemistry, University of Bristol, University Walk, Bristol, BS8 1TD, UK.

**Christopher Williams** – School of Chemistry, University of Bristol, Bristol, BS8 1TS, UK.

**Ethan L. Bungay** – School of Biochemistry, University of Bristol, University Walk, Bristol, BS8 1TD, UK.

**Jessica C. Berrones-Reyes** – Centre for Enzyme Innovation, School of the Environment and Life Sciences, University of Portsmouth, PO1 2DT, UK.

**Rob Barringer** – School of Biochemistry, University of Bristol, University Walk, Bristol, BS8 1TD, UK.

**Catherine R. Back** – School of Biochemistry, University of Bristol, University Walk, Bristol, BS8 1TD, UK.

**Paul Molinaro** – Department of Physics, The City College of New York, New York NY10031; Graduate Programs of Physics, Biology, Chemistry and Biochemistry, The Graduate Center of CUNY, New York, New York 10016, United States

**Ronald L. Koder** – Department of Physics, The City College of New York, New York NY10031;

**Bruce R. Lichtenstein** – Centre for Enzyme Innovation, School of the Environment and Life Sciences, University of Portsmouth, PO1 2DT, UK.

**Adrian J. Mulholland** – School of Chemistry, University of Bristol, Bristol, BS8 1TS, UK.

**Matthew P. Crump** – School of Chemistry, University of Bristol, Bristol, BS8 1TS, UK.

## Author Contributions

The manuscript was written with contributions from all authors. All authors have given final approval to the final version of the manuscript.

## Acknowledgments

This work was supported at the University of Bristol by the Biological and Biotechnological Sciences Research Council (BB/W003449/1 and BB/T008741/1; the latter providing a studentship to C.M.). We also wish to thank the Diamond Light Source staff, particularly those at the i04 beamline.

