## Supporting Information for "Porphyrin-driven redox tuning in structurally defined *de novo* heme proteins"

**Contents.**

**P2-9: Materials and Methods**
**P7: Scheme S1** Synthetic route from hemin to Fe(III) 2,4-dimethyl-deuteroporphyrin IX.
**P10: Table S1** UV/visible spectroscopic data for m4D2:NNP and 4D2 T19D:NNP complexes.
**P11: Table S2** NNP binding affinities to m4D2 and 4D2 T19D.
**P12: Table S3** Crystallographic data collection and refinement statistics for 4D2 T19D:mesoheme and 4D2 T19D:DF-heme structures.
**P13: Table S4** Redox potentials of m4D2:NNP and 4D2 T19D:NNP complexes measured by redox potentiometry.
**P14: Figure S1** Biophysical analysis of m4D2 T19D:heme B.
**P15: Figure S2** Crystallographic analysis of 4D2 T19D:NNP/heme variants.
**P16: Figure S3** 2D NMR analysis of NNP binding to m4D2.
**P17: Figure S4** $2F_O$-$F_C$ OMIT maps reveal variation in NNP orientational preferences in 4D2 T19D.
**P18: Figure S5** Rationalising orientational preference of DF-heme in 4D2 T19D.
**P19: Figure S6** NMR structural ensemble of m4D2:DM-heme and comparison with computational structural predictions.
**P20: Figure S7** HPLC analysis of 2,4-Diformyl-deuteroporphyrin IX dimethyl ester *p*-ditosylhydrazone (4).
**P21: SI References**

### Materials and Methods

#### General

Unless specified here, all chemicals and reagents were purchased from Merck, Thermo Fisher, TCI and VWR. Hemin was purchased from Fluka, Fe(III) deuteroporphyrin IX (sc-396884) and Fe(III) mesoporphyrin IX (sc-396889) from ChemCruz, and Fe(III) 2,4-diacetyldeuteroporphyrin IX (D40050) and Fe(III) 2,4-diformyldeuteroporphyrin IX (D14624) from Frontier Specialty Chemicals. UV/visible spectra were recording using an Agilent Cary 60 spectrometer.

#### Protein expression and purification

Starter cultures of *E. coli* T7 Express (NEB) transformed with m4D2 or 4D2 T19D[1] expression plasmids were grown overnight in 100 mL of LB medium supplemented with 50 µg/mL carbenicillin in a shaking incubator set to 37°C and 180 rpm. These were subsequently used to inoculate 1 L cultures of LB media (50 µg/mL carbenicillin) and grown under the same conditions until $OD_{600} \geq 0.6$. Protein expression was induced by adding 1 mM IPTG and 25 mg/mL δ-aminolevulinic acid (ALA) to enhance intracellular heme loading, followed by incubation for 3–4 h. Cells were harvested by centrifugation at 8,000 x g for 25 min at 4°C. Pellets were resuspended in lysis buffer (50 mM $NaH_2PO_4$, 100 mM NaCl, 20 mM imidazole, pH 8.0) and stored at –20°C prior to lysis and purification.

Cell pellets were thawed and lysed either by sonication on ice (Fisherbrand 120 probe-tip sonicator (Thermo Fisher); 80% amplitude, 20 s on/20 s off cycles for 4 min) or by high-pressure homogenisation using a Constant Systems CF1 Cell Disruptor (25 kpsi) following manual homogenisation. Lysates were clarified by centrifugation at 38,758 x g for 35 min at 4°C, and the supernatant was collected. The supernatant was then applied to a 5 mL HisTrap nickel-affinity column (Cytiva) equilibrated with 50 mL lysis buffer. Bound protein was eluted with elution buffer (50 mM $NaH_2PO_4$, 100 mM NaCl, 250 mM imidazole, pH 8.0). Eluted protein was exchanged into 50 mM Tris-HCl, 0.5 mM EDTA, pH 8.0 for overnight cleavage by tobacco etch virus NIa (TEV) protease (200–400 µg per 2 L of original culture) with the addition of tris(2-carboxyethyl(phosphine) (TCEP) to 1 mM. Uncleaved protein was removed by reverse nickel-affinity chromatography. Cleaved protein was concentrated to ~4.5 mL using 3–5 kDa centrifugal concentrators (VivaSpin) and subjected to size-exclusion chromatography (SEC) on a 120 mL HiLoad 16/600 Superdex 75 pg column (Cytiva) equilibrated with 20 mM CHES, 100 mM KCl, pH 8.6.

#### Heme extraction and NNP loading

Endogenously loaded heme B was removed through acid-butanone heme extraction of purified protein[2]. Heme-containing protein in 20 mM CHES, 100 mM KCl, pH 8.6 was acidified to pH 2.0 through dropwise addition of concentrated HCl and then mixed with an equivalent volume of methyl ethyl ketone (butanone) on ice. This mixture was applied to a separating funnel and further agitated, then allowed to separate into organic and aqueous phases. The lower protein-containing layer was collected, and the procedure was repeated twice to ensure complete heme removal. Heme-extracted apoproteins were dialysed against 4 L of 1 mM $NaHCO_3$, then 2 x 4 L of 20 mM CHES, 100 mM KCl, pH 8.6, using SnakeSkin dialysis tubing (Thermo Fisher) with 3.5 kDa cut off. NNP stocks of 1-10 mg/mL were prepared by dissolution in dimethyl sulfoxide and stored at -20 °C. To load apoproteins with these heme analogues, NNP solutions were added to the apoproteins at 1.5x molar excess (based on the number of heme binding sites in the respective protein) and incubated at room temperature for at least 30 minutes prior to application to a PD-10 column (Cytiva) loaded with Sephadex G-25 (Merck) to remove excess NNP.

**Protein concentration determination**

Apoprotein extinction coefficients at 280 nm ($\varepsilon_{280nm}$) were calculated using the ProtParam tool[3] on the ExPASy website[4], and concentrations were determined using UV/visible spectroscopy. Protein concentrations were confirmed using the Bradford assay; briefly, a calibration curve was created using Bradford's reagent (VWR) and stock solutions of bovine serum albumin (BSA, VWR) according to the manufacturer's protocol. Apo- or holoprotein concentration was determined by adding 100 µL of sample to 1 mL reagent and comparing to the BSA calibration. All measurements were recorded in triplicate.

**Pyridine hemochromogen assay**

The pyridine hemochromogen assay[5] was used to determine the extinction coefficient (ε) of free and protein-bound porphyrins as previously described. 20 µL of protein sample (50 – 100 µM) was added to 100 µL of buffer, 15 µL of 5 M NaOH and 15 µL of pyridine, and a visible spectrum of this mix was recorded, representing the oxidised pyridine hemochromogen. Several grains of sodium dithionite were subsequently added, and visible spectra were recorded until the fully reduced pyridine hemochromogen spectrum was obtained. For heme B, and the NNPs used here, published extinction coefficients could then be used to determine their concentration in the protein sample.

**Isotopic labelling for NMR**

To enable the acquisition of high quality, multidimensional NMR spectra, m4D2 T19D was $^{15}$N-labelled as previously described[1]. Briefly, 100 mL LB cultures of m4D2 T19D-expressing *E. coli* were grown overnight as described above. These were used to inoculate 1 L LB cultures which were grown until $OD_{600nm}$ reached 0.6, after which the cells were harvested by centrifugation (5,000 x g for 10 minutes) in ethanol-sterilised centrifuge flasks. The pelleted cells were then resuspended in 200 mL of M9 salt solution and re-pelleted by centrifugation; this was repeated once more, and the resulting pellet was resuspended in 750 mL of M9 minimal media containing $^{15}NH_4Cl$. These cells were incubated for 1 hour at 37 °C and 180 rpm to enable the cells to recover from the previous wash steps and then induced with 1 mM IPTG. Protein was then expressed for four hours at 37°C and 180 rpm, then harvested and purified as described above.

**Heme B and NNP binding titrations**

Porphyrin binding affinities were determined through titrations monitored by UV/visible spectroscopy. Apoprotein solutions (1 mL at 1-3 µM) were prepared in 20 mM CHES, 100 mM KCl, pH 8.6, and small volumes (1-2 µL) of NNP stocks in DMSO (1-2 mg/mL) were sequentially added. After each addition, equilibration was monitored using UV/visible spectroscopy, with a final, saved absorbance spectrum representing each fully equilibrated NNP concentration. Data were collected in triplicate, averaged and fitted to the tight binding equation (1) to determine the dissociation constant, $K_D$:

$$Absorbance = [C] \times \varepsilon_{free} + \left(\varepsilon_{bound} - \varepsilon_{free}\right) \frac{K_D + [P_{tot}] + [C] - \sqrt{(K_D + [P_{tot}] + [C])^2 - 4[P_{tot}][C]}}{2} \quad (1)$$

*Where:*
[C] = Cofactor Concentration (μM)
[$P_{tot}$] = Total Protein Concentration (μM)
$\varepsilon_{free}$ = Unbound Cofactor Extinction Coefficient ($\mu M^{-1} cm^{-1}$)
$\varepsilon_{bound}$ = Bound Cofactor Extinction Coefficient ($\mu M^{-1} cm^{-1}$)

**Stopped-flow spectrophotometry**
The kinetics of porphyrin binding were determined using a SX20 Stopped Flow Spectrophotometer (Applied Photophysics) equipped with a diode-array detector. Stoichiometric quantities of apoprotein (10 μM) and NNP (10 μM), each in 20 mM CHES, 100 mM KCl, pH 8.6, were rapidly mixed at 22 °C, and UV/visible spectra (250 – 800 nm) were recorded. Single wavelength kinetic data was subsequently extracted for analysis.

**Circular Dichroism (CD) spectroscopy**
Protein secondary structure and thermal stability were determined using a J-1500 Circular Dichroism spectrometer (JASCO). CD spectra were recorded in a 1 mm pathlength quartz cuvette and were collected in 1 °C increments from 5 – 95 °C for the melt, then from 95 – 5 °C to monitor the refold. Raw CD data was converted into Mean Residue Ellipticity (MRE) using equation 2:

$$MRE\ (deg.cm^2.dmol^{-1}) = \frac{Raw\ ellipticity\ (mdeg) \times 10^6}{Pathlength\ (mm) \times [Protein](\mu M) \times n} \quad (2)$$

Where possible, a two-state Boltzmann model (3) was applied to the single wavelength melt data to determine the melting temperature, $T_m$:

$$CD(T) = CD_{min} + \frac{CD_{max} - CD_{min}}{1 + e^{(Tm - T)/s}} \quad (3)$$

*Where:*
*CD(T)* = CD signal at 222 nm at temperature *T.*
$CD_{min}$ = Maximum CD signal at 222 nm
$CD_{max}$ = Minimum CD signal at 222 nm
$T_m$ = Melting temperature
*s* = Gradient of the unfolding transition

**Redox potentiometry**
Heme midpoint potentials were determined using Optically Transparent Thin Layer Electrochemistry (OTTLE) as previously described[1, 6]. Briefly, NNP- and heme-bound proteins (~50 μM) were prepared in 20 mM CHES, 100 mM KCl, pH 8.6 and 10 % glycerol, and 1 μL of redox mediators (1-2 mg/mL) were added to cover the relevant

potential range scanned. These included anthroquinone-sulfonate, phenazine, 2-hydroxy-1,4-naphthoquinone, duroquinone, indigotrisulfonate, phenazine ethosulfate, phenazine methosulfate, *N,N,N',N'*-tetramethyl-*p*-phenylene diamine, and potassium ferricyanide. Potential was applied across a custom quartz electrochemical cell using a Biologic SP-150 potentiostat, platinum working and counter electrodes, and a Ag/AgCl reference electrode (RE-5B, BASi). UV/visible spectra were recorded at each applied potential and after 30 minutes of equilibration. Potentials were converted to mV vs the Nernst Hydrogen Electrode (NHE), using calibration datasets against cytochrome *c*. Midpoint redox potentials were determined by fitting plotted data (applied potential vs heme/NPP ferrous Soret absorbance) using either one (4) or two sequential (5) Nernst equations. All data were recorded in triplicate.

$$1e^{-} \qquad Abs = \left( A + B.10^{\frac{mV-E_m}{59}} \right) / \left( 1 + 10^{\frac{mV-E_m}{59}} \right) \tag{4}$$

$$2 \times 1e^{-} \qquad Abs = \left( A.10^{\frac{mV-E_{m1}}{59}} + C + B.10^{\frac{mV-E_{m2}}{59}} \right) / \left( 1 + 10^{\frac{mV-E_{m1}}{59}} + 10^{\frac{mV-E_{m2}}{59}} \right) \tag{5}$$

**X-ray crystallography**

Crystals of 4D2 T19D loaded with NNPs were grown using a condition adapted from previous work. Purified proteins in 20 mM CHES, 100 mM KCl at pH 8.6 were concentrated to concentration of 5 – 15 mg/mL$^{-1}$ and mixed with 2.1 M DL-Malic acid, pH 7.0 at 1:1 and 3:1 protein:precipitant ratios, in 1 µL hanging droplets. Trays were left at 20 °C and produced well-formed crystals within a month. Crystals were mounted in LithoLoops (Molecular Dimensions), flash-frozen in liquid nitrogen either without cryoprotectant (4D2 T19D:mesoheme) or dipped in ethylene glycol (4D2 T19D::DF-heme) and sent to the i04 beamline at Diamond Light Source, UK for illumination. Diffraction data preocessing, reduction and scaling were performed using Xia2 multiplex[7] and the AutoPROC[8] for Mesoheme and DF-Heme loaded 4D2 T19D crystals, respectively. Molecular replacement was performed using Molrep[9] implementing the 4D2 T19D crystal structure for the Mesoheme form (PDB ID: 8CCR) and the predicted AlphaFold[10] structure for the DF-heme form. Models were refined iteratively using Coot[11] and Refmac[12] within the CCP4i2[13] software suite.

**NMR spectroscopy**

For structure studies, 0.5 -1 mM protein was dissolved in 20 mM $KH_2PO_4$, 50 mM KCl pH 6.4 with 10% $D_2O$. NMR experiments were acquired at 25 °C or 35 °C on a 700 MHz Bruker *AVANCE HD III* spectrometer equipped with a 1.7 mm TCI micro-cryoprobe or a 800 MHz Bruker *AVANCE III HD* spectrometer equipped with a 5 mm TCI cryoprobe (New York Structural Biology Center, NYSBC). The backbone resonances were assigned using HNCA/HNCOCA/HNCACB/HNCOCACB and HNCO/HNCACO spectra. Side-chain resonances were assigned using CCONH, $^{15}$N-edited TOCSY, $^{15}$N-edited NOESY, HCCH-TOCSY and $^{13}$C-edited NOESY spectra. NMR data were processed in NMRPipe v11[14] and analyzed using CCPNMR analysis v2.4.2[15]. $^{1}$H-$^{15}$N TROSY-HSQC experiments probing NPP binding were performed on a 700 MHz Bruker Avance III HD spectrometer equipped with a 1.7 mm TCI micro-cryoprobe at 298K using the standard best TROSY pulse sequence from the Bruker library.

Structure calculations were performed using ARIA v2.3.2[16] coupled to CNS v1.21[17] using NOE distance restraints and dihedral restraints calculated from $^{1}H_{\alpha}$, $^{15}N$ and $^{13}C_{\alpha}/C_{\beta}$ chemical shifts using TALOS[18]. Spin diffusion was enabled throughout, and default parameters were used, apart from the number of steps, which were increased for all parts of the calculation to 25000 high-temperature steps, 80000 cool1 and cool2 steps. Hydrogen bond restraints identified from the preliminary rounds of calculations and from the two-heme binding 4D2 homologue (7AH0) along with iron 2,4-dimethyl-deuteroporphyrin IX (DM-heme; FDD in the PDB and BMRB) were introduced at the later stages of refinement once the protein had converged. Due to bleaching out of the NMR signals close to the metal center, only a few tentative heme-protein intermolecular NOEs could be identified in the NOESY spectra. These were included in the calculation as ambiguous NOEs. However, due to the symmetry of the DM-heme, these were not sufficient to fully fix the orientation of the heme in the binding site. Therefore, a small number of unambiguous restraints derived from the two-heme binding 4D2 homologue (7AH0) were also used to restrain the orientation of the heme during the calculation. For the final calculation, 500 structures were calculated in iteration 8, of which the fifty lowest energy structures were further water refined. Twenty structures with the lowest energy and least number of violations were selected for the final ensemble. PSVS v2.0[19] was used to validate the structures. The chemical shifts and structural restraints were deposited with the Biological Magnetic Resonance Data Bank (BMRB) under accession number 34977. The structure was deposited with the PDB under code 9I38.

**Synthetic route to iron(III) 2,4-dimethyldeuteroporphyrin IX**

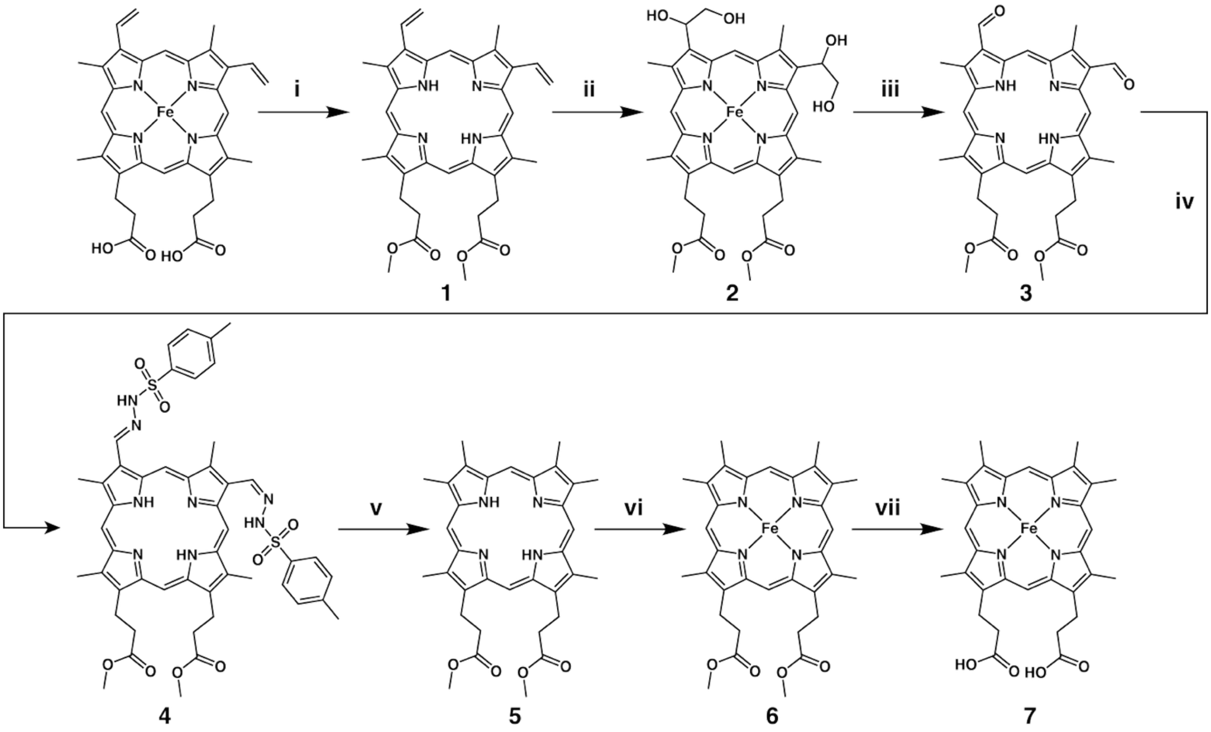


**Scheme S1. Synthetic route from hemin to Fe(III) 2,4-dimethyldeuteroporphyrin IX.** Conditions: **(i)** Pyridine, $FeSO_4$, $SOCl_2$, methanol; **(ii)** $K_2OsO_4$, NMO, dioxane-$H_2O$; **(iii)** periodic acid, $H_2O$, THF; **(iv)** tosylhydrazine, methanol; **(v)** $Na[BH_3(CN)]$, DMF; **(vi)** $FeCl_2.4H_2O$, MeCN, $CHCl_3$; **(vii)** methanol, $H_2O$, KOH.

Protoporphyrin IX dimethyl ester **(1)**

Hemin (2.5 g, 3.84 mmol) was dissolved in 10 mL of pyridine and then diluted to a total volume of 150 mL with methanol. Ferrous sulphate (6.0 g, 21.6 mmol) was added to the solution. While stirring, 25 mL of thionyl chloride (344 mmol) was introduced using a dropping funnel, added at a rate fast enough to induce reflux in the reaction mixture without the need for external heating. The reaction vessel was kept open to the atmosphere during the addition of thionyl chloride to prevent over-pressurization, which could lead to the vessel shattering. Once the thionyl chloride addition was complete, the reaction mixture was stirred under an argon atmosphere for 30 minutes, allowing it to cool to room temperature. Argon was then bubbled through the solution for an additional 30 minutes to remove excess HCl and other volatile acids. It is crucial to complete the workup as quickly as possible, as the free base is sensitive to aqueous acid. The reaction was quenched with 25 mL of water and then diluted with 300 mL of chloroform. The chloroform layer was washed with 300 mL of water, and the aqueous layer was extracted three times with 100 mL portions of chloroform. The combined organic layers were washed six times with 300 mL of water, followed by neutralisation with two washes of 300 mL 1 N $NH_4OH$. The organic phase was dried over magnesium sulphate, and solvents were removed under reduced pressure (2.08 g, yield 92%). UV-vis (DMSO) **λ** max: 402, q bands: 500, 533, and 570 nm. HRMS (MALDI) calculated for $C_{36}H_{38}N_4O_4$ [M+H]+: 591.2966, found: 591.2977.

2,4-Bis(1,2-dihydroxyethyl)-deuteroporphyrin IX dimethyl ester **(2)**[20]

Protoporphyrin IX dimethyl ester **1** (3.38 g, 5.72 mmol) was dissolved in a mixture of dioxane (900 mL) and water (100 mL), which had been vigorously purged with argon. $K_2OsO_4$ (473 g, 1.28 mmol) and 4-methylmorpholine N-oxide (2.13 g, 18.8 mmol) were then added to the dioxane solution. The reaction was stirred in the dark under an argon atmosphere for 24 hours. Sodium metabisulfite (22 g, 115 mmol) was added to the reaction mixture in one portion at room temperature. After stirring for 1 hour at room temperature, the mixture was gradually heated to 65 °C. The temperature was maintained for 15 minutes, after which the solution was allowed to cool to 53 °C. The reaction mixture was then filtered, and the resulting solid was washed with 1,4-dioxane until the filtrate was colourless. The filtrate was reduced to 100 mL under vacuum and slow addition of water allowed the crystallisation of the porphyrin. The product was filtered, rinsed with water, and dried under vacuum. Recrystallization using a mixture of 10% MeOH/$CH_2Cl_2$ (450 mL) and hexanes (550 mL) yielded the desired bis-glycol product (1.8 g, 48% yield). UV-vis (DMSO) **λ** max: 402, q bands: 500, 533, and 570 nm. HRMS (MALDI) calculated for $C_{36}H_{40}N_4O_8$ [M+H]+: 657.2919, found: 657.2926.

2,4-Diformyl-deuteroporphyrin IX dimethyl ester **(3)**[20]

Bis-glycol **2** (300 mg, 0.45 mmol) was suspended in THF (100 mL). To this solution, periodic acid (650 mg, 2.85 mmol) dissolved in water (12 mL) was added, and the mixture was stirred for 1 hour. The solution was then concentrated under reduced pressure, diluted with chloroform, and washed with water. After evaporating the solvent, the crude porphyrin was dissolved in hot $CHCl_3$ and precipitated with hexane, yielding 262 mg (98%). UV-vis (DMSO) **λ** max: 430, q bands: 520, 555, and 588 nm. HRMS (MALDI) calculated for $C_{34}H_{34}N_4O_6$ [M+H]+: 595.2551, found: 595.2559.

2,4-Diformyl-deuteroporphyrin IX dimethyl ester *p*-ditosylhydrazone **(4)**

Diformyl product **3** (2 g, 3.36 mmol) was added to a solution of tosylhydrazine (890 mg, 4.71 mmol) in methanol (20 mL), and the mixture was stirred at 85 °C for 4 h. After cooling to ambient temperature, the solvent was evaporated, and the product was purified by recrystallization from methanol (2.29 g, 73% yield). UV-vis (DMSO) **λ** max: 424, q bands: 518, 553, and 586 nm. HRMS (MALDI) calculated for $C_{48}H_{50}N_8O_8S_2$ [M+H]+: 931.3266, found: 931.3277. HPLC analysis can be found in **Figure S7**.

2,4-Dimethyl-deuteroporphyrin IX dimethyl ester **(5)**

To a solution of compound **4** (930 mg, 1 mmol) in DMF (10 mL), Na[$BH_3$(CN)] (251 mg, 4 mmol) was added, and the mixture was refluxed overnight. The solvent was removed under reduced pressure, and the residue was dissolved in dichloromethane (100 mL) and washed with water. The combined organic layers were dried over $Na_2SO_4$, filtered, and concentrated. The pure product was obtained after several rounds of recrystallization from dichloromethane and hexane (298 mg, 53% yield). UV-vis (DMSO) **λ** max: 402, q bands: 500, 533, and 570 nm. HRMS (MALDI) calculated for $C_{34}H_{38}N_4O_4$ [M+H]+: 567.2966, found: 567.2975.

Iron(III) 2,4-Dimethyl-deuteroporphyrin IX dimethyl ester **(6)**[21]

Ferrous chloride hydrate (110 mg, 0.55 mmol) was placed into a three-necked round-bottom flask equipped with a reflux condenser (connected to an argon inlet), a dropping funnel (fitted with a septum cap), and a second septum cap. The flask was evacuated and refilled with argon three times. Acetonitrile (20 mL), previously refluxed under argon with vigorous stirring for 1 hour, was transferred to the flask via cannula, and the mixture was stirred. Once the ferrous chloride was dissolved, the mixture was cooled to room temperature. In a separate step, 2,4-Dimethyl-deuteroporphyrin IX dimethyl ester (**5**) (80 mg, 0.14 mmol) was dissolved in argon-purged chloroform (10 mL) and introduced through the septum-capped dropping funnel. The porphyrin solution was added dropwise to the stirred ferrous chloride solution at a rate of 2 mL per minute. After the addition was complete, the mixture was stirred under an argon atmosphere for 20 minutes before being exposed to air for 10 minutes. The brown solution was then diluted with dichloromethane (20 mL), followed by successive washes with 0.2 M HCl and water. The organic layer was dried over $Na_2SO_4$, and the solvent was evaporated. Recrystallization of the residue from dichloromethane and heptane yielded 82 mg of product (95% yield). UV-vis (DMSO) **λ** max: 395. HRMS (ESI) calculated for $C_{34}H_{36}N_4O_4Fe$[M+H]+: 620.2086, found: 620.2071.

Iron(III) 2,4-Dimethyl-deuteroporphyrin IX **(7)**[21]

Iron(III) 2,4-Dimethyl-deuteroporphyrin IX dimethyl ester (**6**) (50 mg, 0.08 mmol) was dissolved in 15 mL of a solution prepared by mixing 95 mL of methanol, 5 mL of water, and 1 g of KOH. The resulting solution was refluxed under an argon atmosphere overnight. The mixture was then diluted with dichloromethane (20 mL) and washed twice with 0.2 M HCl (50 mL). The organic phase was then collected, dried ($Na_2SO_4$), and evaporated to dryness yielding a residue that was recrystallized from a THF-hexane mixture (36 mg, 75% yield). UV-vis (DMSO) **λ** max: 395. HRMS (ESI) calculated for $C_{32}H_{32}N_4O_4Fe$ [M+H]+: 592.1773, found: 592.1786.

| m4D2 (nm) | | | | | | | | |
|---|---|---|---|---|---|---|---|---|
| ***Peak*** | ***Deuteroheme*** | | ***Mesoheme*** | | ***DA-heme*** | | ***DF-heme*** | |
| | *ferric* | *ferrous* | *ferric* | *ferrous* | *ferric* | *ferrous* | *ferric* | *ferrous* |
| *γ* | 403 | 413 | 406 | 416 | 430 | 454 | 438 | 464 |
| *β* | 514 | 520 | 520 | 523 | 547 | 549 | 555 | 553 |
| *α* | 552 | 548 | 553 | 551 | 586 | 582 | 595 | 594 |
| **4D2 T19D (nm)** | | | | | | | | |
| | ***Deuteroheme*** | | ***Mesoheme*** | | ***DA-heme*** | | ***DF-heme*** | |
| | *ferric* | *ferrous* | *ferric* | *ferrous* | *ferric* | *ferrous* | *ferric* | *ferrous* |
| *γ* | 404 | 412 | 406 | 415 | 430 | 455 | 440 | 464 |
| *β* | 520 | 519 | 520 | 522 | 545 | 549 | 556 | 554 |
| *α* | 554 | 548 | 552 | 551 | 586 | 583 | 596 | 593 |

**Table S1. UV/visible spectroscopic data for m4D2:NNP and 4D2 T19D:NNP complexes.** All peak positions are provided in nanometres (nm) and recorded on samples at 2-10 μM in 20 mM CHES, 100 mM KCl, pH 8.6.

| ***Protein*** | ***NNP*** | ***$K_D$ (nM)*** | ***SD (nM)*** |
|---|---|---|---|
| **m4D2** | *Deuteroheme* | 1.45 | 1.1 |
| | *Mesoheme* | 0.84 | 1.7 |
| | *DA-heme* | 15.6 | 7.5 |
| | *DF-heme* | 12.0 | 7.7 |
| **4D2 T19D** | *Deuteroheme* | 30.3 | 9.3 |
| | *Mesoheme* | 18.3 | 5.1 |
| | *DA-heme* | 30.4 | 7.5 |
| | *DF-heme* | 8.32 | 3.0 |
| | *Heme B* | 20.0 | 2.9 |

**Table S2. NNP binding affinities to m4D2 and 4D2 T19D.** All data was measured in triplicate, with $K_D$ *representing the average binding constant. SD indicates the corresponding standard deviation*. All measurements were recorded on samples at 1-3 μM in 20 mM CHES, 100 mM KCl, pH 8.6.

| Crystallography data collection and refinement | | |
|---|---|---|
| Protein | 4D2 T19D:Mesoheme | 4D2 T19D:DF-heme |
| PDB ID | 9H4C | 9SY9 |
| **Data collection** | | |
| Space group | H3 | H3 |
| Unit cell dimensions (Å): A, B, C | 81.19, 81.19, 58.82 | 81.27, 81.27, 60.97 |
| Unit cell angles (°): α, β, γ | 90, 90, 120 | 90, 90, 120 |
| Resolution (Å) | 45.11 - 1.98 (2.01 - 1.98) | 46.08 - 2.00 (2.03 - 2.00) |
| CC1/2 (%) | 0.99 (0.40) | 1.00 (0.52) |
| $R_{pim}$ | 0.08 (-11.318) | 0.03 (0.74) |
| Number of unique reflections | 10079 (504) | 10188 (513) |
| Multiplicity | 85.6 (87.2) | 10.7 (9.2) |
| Overall signal-to-noise ratio (I/σ) | 11.2 (0.2) | 11.8 (1.1) |
| Completeness (%) | 100 (100) | 100 (100) |
| **Refinement** | | |
| Rwork | 0.20 (0.46) | 0.18 (0.38) |
| Rfree | 0.25 (0.51) | 0.23 (0.34) |
| No protein atoms used in refinement | 920 | 865 |
| No water atoms used in refinement | 6 | 11 |
| B factors for protein atoms (Å$^2$) | 59 | 46 |
| B factors for ligand atoms (Å$^2$) | 54 | 46 |
| B factors for water atoms (Å$^2$) | 68 | 58 |
| RMS deviations—length (Å) | 0.67 | 0.60 |
| RMS deviations—angle (°) | 1.48 | 1.29 |
| Ramachandran-favored residues (%) | 99 | 99 |
| Ramachandran outlying residues (%) | 0 | 0 |

**Table S3. Crystallographic data collection and refinement statistics for 4D2 T19D:mesoheme and 4D2 T19D:DF-heme structures.** Values in parenthesis are for the highest resolution shell.

| ***Protein*** | ***NNP*** | ***Em (mV)*** | ***SD (mV)*** |
|---|---|---|---|
| **m4D2** | *Deuteroheme* | -218 | 1.9 |
| | *Mesoheme* | -235 | 1.3 |
| | *DA-heme* | 29.9 | 1.9 |
| | *DF-heme* | 141 | 7.1 |
| **4D2 T19D** | *Deuteroheme* | -171, -232 | 2.1, 2.6 |
| | *Mesoheme* | -188, -257 | 5.6, 3.5 |
| | *DA-heme* | -14.5, 80.4 | 7.7, 1.8 |
| | *DF-heme* | 99.6, 169 | 2.8, 2.2 |

**Table S4. Redox potentials of m4D2:NNP and 4D2 T19D:NNP complexes measured by redox potentiometry.** Potentials were calculated by fitting redox potentiometry data to one (m4D2 data) or sequential two (4D2 T19D) electron Nernst equations. All measurements were recorded on samples prepared in 20 mM CHES, 100 mM KCl, pH 8.6.

### Supplementary Figures

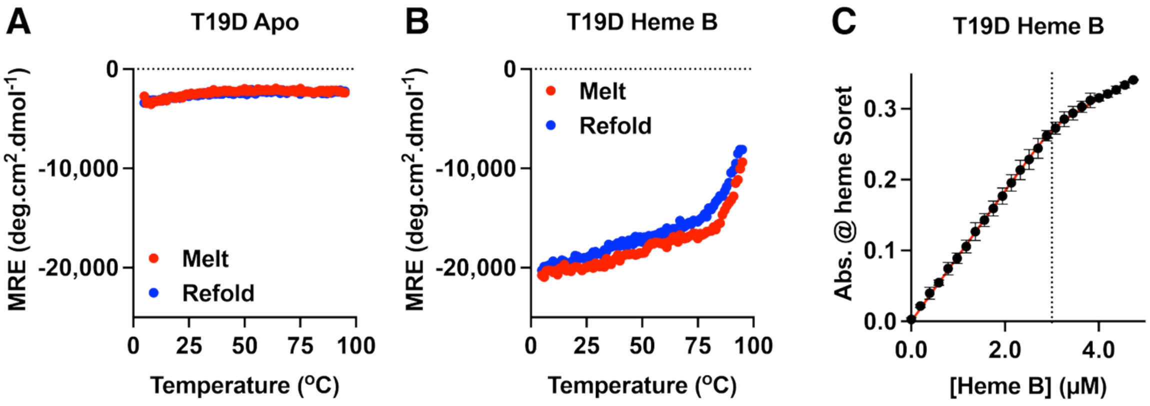


**Figure S1. Biophysical analysis of 4D2 T19D:hemeB.** Temperature-dependent CD signal at 222 nm for apo-4D2 T19D (A) and 4D2 T19D:heme B (B). CD data were collected on 1-3 μM apoprotein in 20 mM CHES, 100 mM KCl, pH 8.6. (C) Heme B binding isotherm for 4D2 T19D, recorded at 1 μM protein in 20 mM CHES, 100 mM KCl, pH 8.6. Error bars represent standard deviations for each point in the binding isotherms, and the dashed line indicates a 2:1 stoichiometric ratio of heme B:protein.

A

B

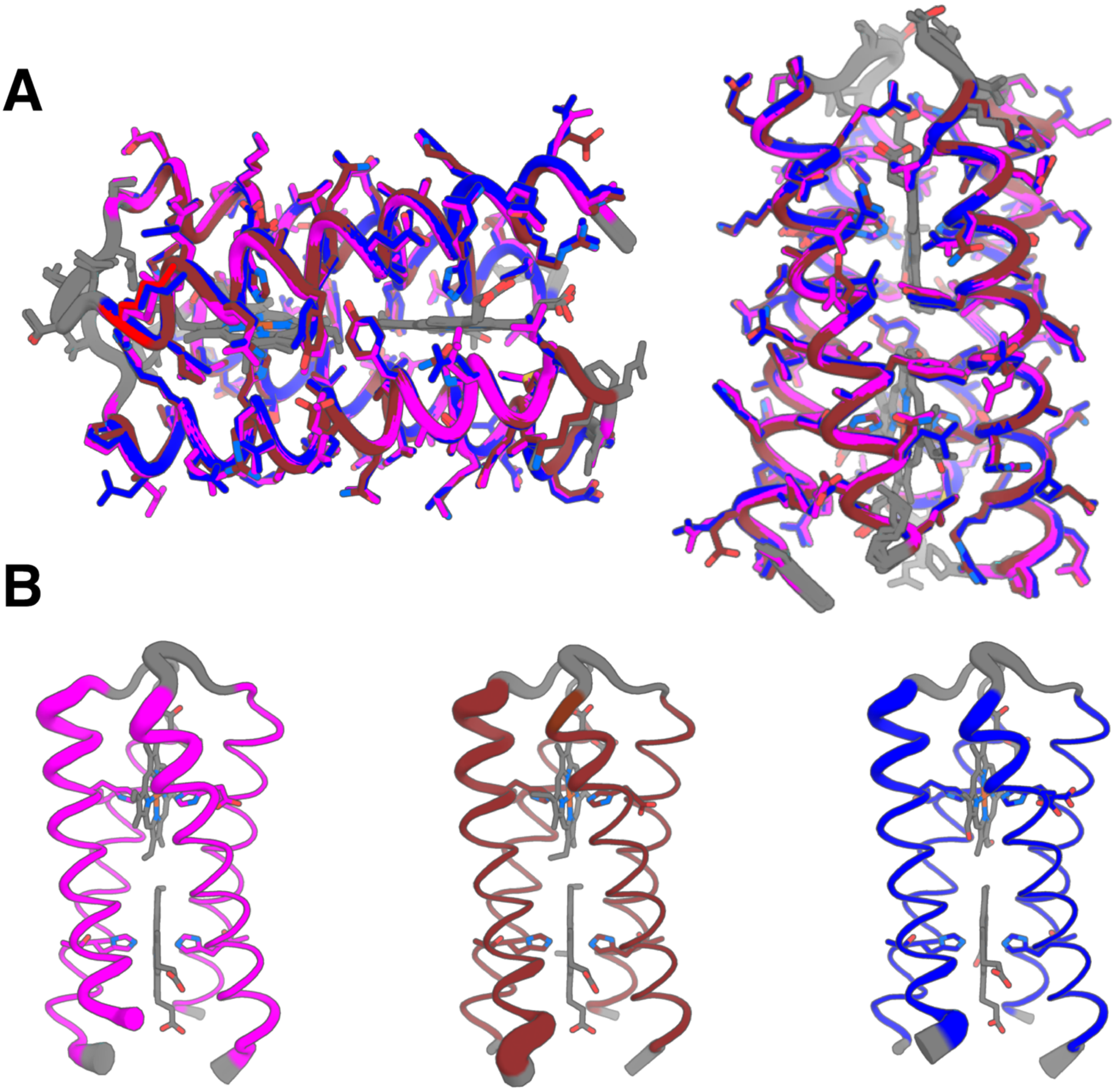

**Figure S2. Crystallographic analysis of 4D2 T19D:NNP/heme variants.** (A) Overlay of 4D2 T19D with heme B (PDB 8CCR; magenta), mesoheme (PDB 9H4C; chocolate) and DF-heme (PDB 9SY9; blue), showing near-identical positioning of sidechains in each variant. (B) Crystallographic B-factors for 4D2 T19D:NNP complexes**.** B-factors are displayed for 4D2 T19D with heme B (PDB 8CCR; magenta, left), mesoheme (PDB 9H4C; chocolate, centre) and DF-heme (PDB 9SY9; blue, right).

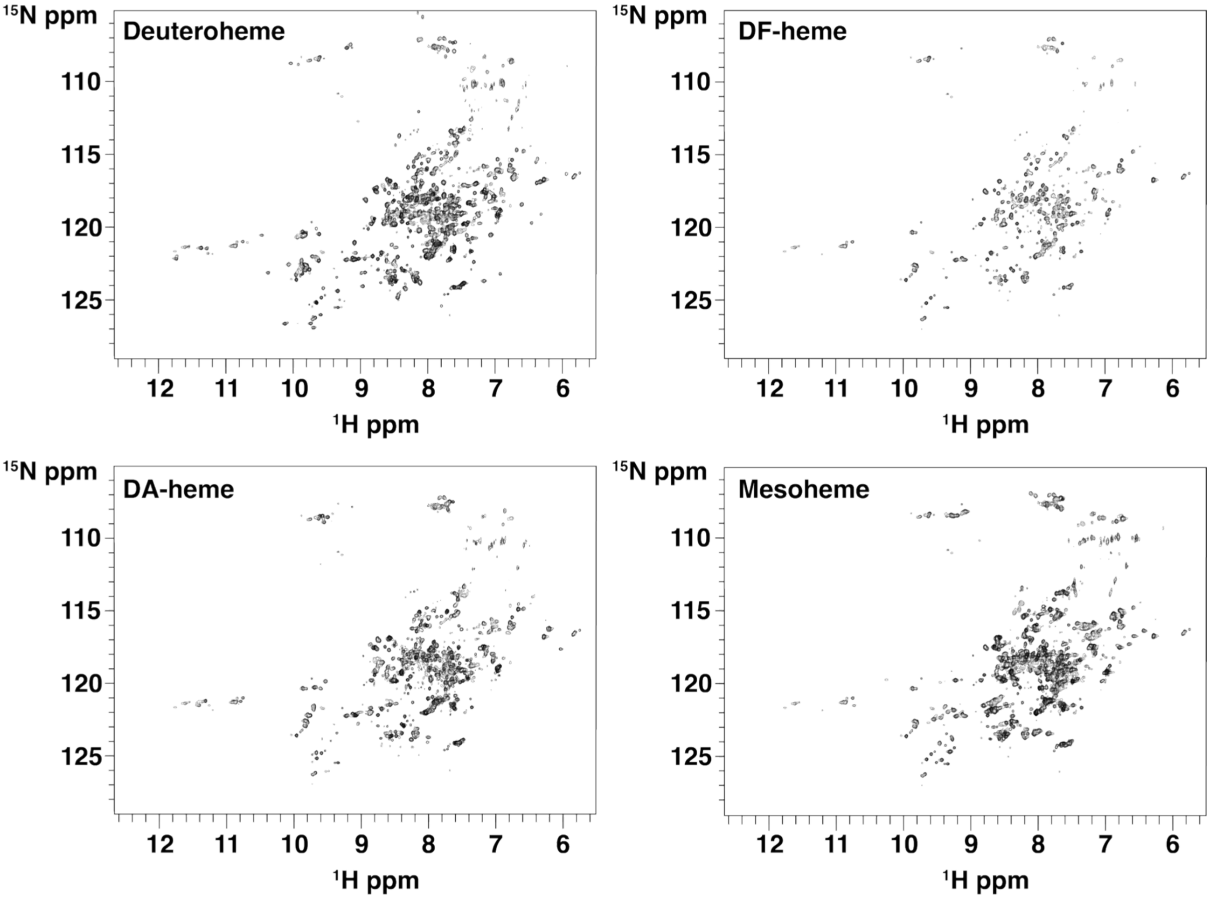


**Figure S3. 2D NMR analysis of NNP binding to m4D2.** $^{15}N$-$^{1}H$ TROSY NMR spectrum of deuteroheme, mesoheme, DF-heme and DA heme binding to m4D2 collected at 700 MHz. Data were recorded on $^{15}N$-labelled protein at 200 μM in 20 mM $KH_2PO_4$, 50 mM KCl, pH 6.4 and 25 °C.

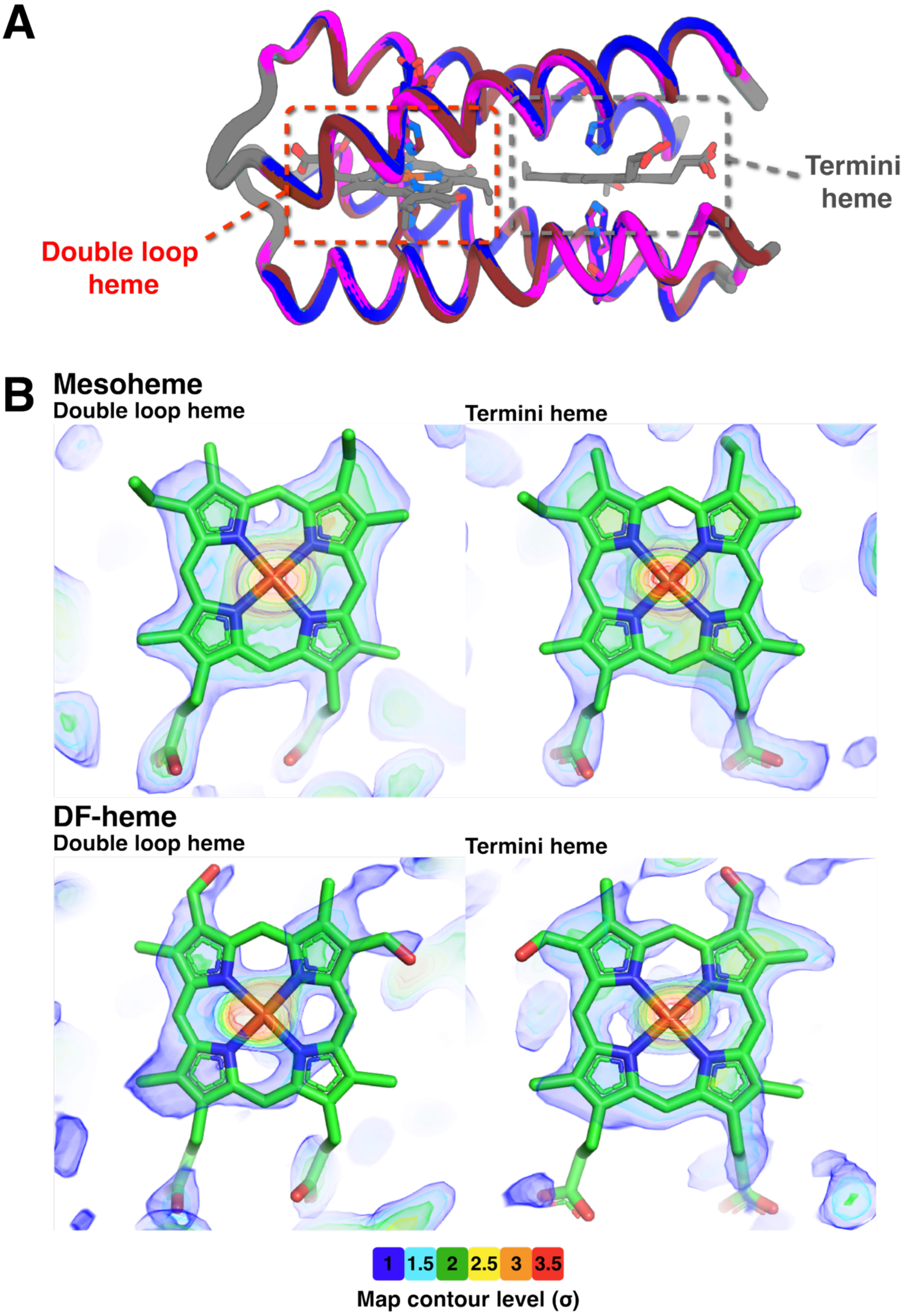


**Figure S4. $2F_O$-$F_C$ OMIT maps reveal variation in NNP orientational preferences in 4D2 T19D.** (A) Definition of heme/NNP binding sites based on their location at the protein end with two loops (double loop heme) or a single loop and the termini (termini heme). (B) $2F_O$-$F_C$ OMIT maps derived from X-ray crystallographic analysis of 4D2 T19D binding mesoheme (PDB 9H4C) and DF-heme (9SY9).

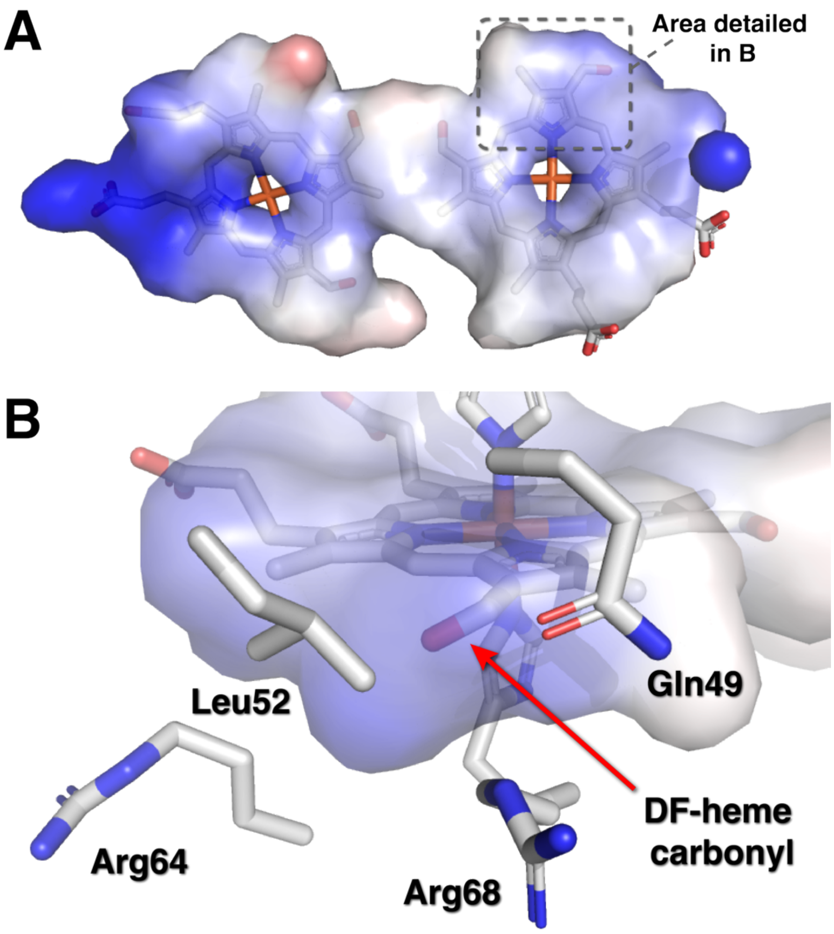


**Figure S5. Rationalising orientational preference of DF-heme in 4D2 T19D.** (A) Electrostatic surface potential map of the heme binding sites in DF-heme-loaded 4D2 T19D. (B) Detail of the termini heme binding site, with amino side chains key to defining the surface electrostatic and hydrophobic properties indicated.

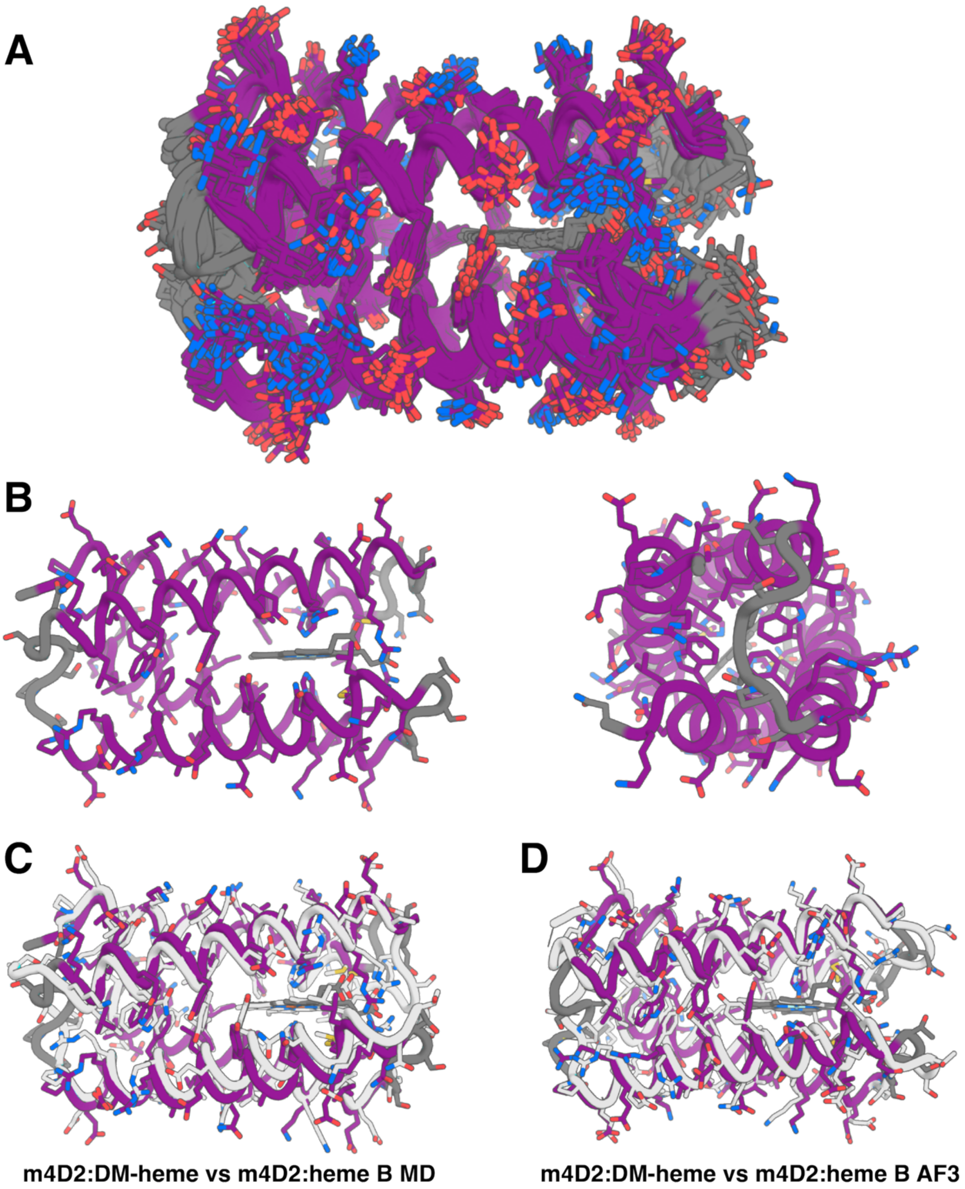


**Figure S6. NMR structural ensemble of m4D2:DM-heme and comparison with computational structural predictions.** (A) 20 lowest energy structures of m4D2:DM-heme from NMR analysis (PDB 9I38). (B) Side and end views of the lowest energy structure from the NMR ensemble. (C) Overlay of the m4D2:DM-heme lowest energy structure with a computational model of m4D2:heme B from the final frame of a molecular dynamics simulation[1]. (D) Overlay of the m4D2:DM-heme lowest energy structure with an AlphaFold3 prediction of the m4D2:heme B complex[22].

### Single Injection Report

| | | | |
|---|---|---|---|
| **Sample name:** | Compound 4 | | |
| **Data file:** | 2024-01-19 19-32-54+00-00-11.dx | **Operator:** | CEI User (CEI User) |
| **Instrument:** | LEFT-HAND SIDE HPLC (DAD - 1200 Series) | **Injection date:** | 2024-01-19 19:33:37+00:00 |
| **Inj. volume:** | 10.000 µL | **Location:** | 5 |
| **Acq. method:** | Luna_95%MetOH-5%Water(0.1%TFA).amx | **Type:** | Sample |
| **Processing method:** | | **Calib Level:** | |
| | | **Sample amount:** | 0.00 |
| **Manually modified:** | None | | |

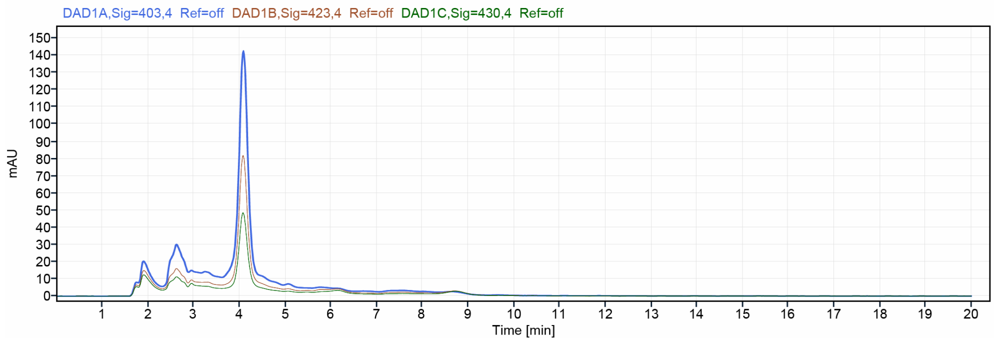




**Figure S7. HPLC analysis of 2,4-Diformyl-deuteroporphyrin IX dimethyl ester *p*-ditosylhydrazone (4).** Analysis was performed on a Luna C18, 5 µm column (150 × 4.6 mm) using an isocratic mobile phase of 95% methanol/5% water containing 0.1% TFA at a flow rate of 0.6 mL min$^{-1}$, with UV detection at 403 nm.

(13) Potterton, L.; Agirre, J.; Ballard, C.; Cowtan, K.; Dodson, E.; Evans, P. R.; Jenkins, H. T.; Keegan, R.; Krissinel, E.; Stevenson, K.; et al. CCP4i2: the new graphical user

interface to the CCP4 program suite. *Acta Crystallogr D Struct Biol* **2018**, *74* (Pt 2), 68–84. DOI: 10.1107/S2059798317016035 From NLM Medline.
(14) Delaglio, F.; Grzesiek, S.; Vuister, G. W.; Zhu, G.; Pfeifer, J.; Bax, A. NMRPipe: a multidimensional spectral processing system based on UNIX pipes. *J Biomol NMR* **1995**, *6* (3), 277–293. DOI: 10.1007/BF00197809 From NLM Medline.
(15) Vranken, W. F.; Boucher, W.; Stevens, T. J.; Fogh, R. H.; Pajon, A.; Llinas, M.; Ulrich, E. L.; Markley, J. L.; Ionides, J.; Laue, E. D. The CCPN data model for NMR spectroscopy: development of a software pipeline. *Proteins* **2005**, *59* (4), 687–696. DOI: 10.1002/prot.20449 From NLM Medline.
(16) Rieping, W.; Habeck, M.; Bardiaux, B.; Bernard, A.; Malliavin, T. E.; Nilges, M. ARIA2: automated NOE assignment and data integration in NMR structure calculation. *Bioinformatics* **2007**, *23* (3), 381–382. DOI: 10.1093/bioinformatics/btl589 From NLM Medline.
(17) Brunger, A. T. Version 1.2 of the Crystallography and NMR system. *Nat Protoc* **2007**, *2* (11), 2728–2733. DOI: 10.1038/nprot.2007.406 From NLM Medline.
(18) Shen, Y.; Delaglio, F.; Cornilescu, G.; Bax, A. TALOS+: a hybrid method for predicting protein backbone torsion angles from NMR chemical shifts. *J Biomol NMR* **2009**, *44* (4), 213–223. DOI: 10.1007/s10858-009-9333-z From NLM Medline.
(19) Bhattacharya, A.; Tejero, R.; Montelione, G. T. Evaluating protein structures determined by structural genomics consortia. *Proteins* **2007**, *66* (4), 778–795. DOI: 10.1002/prot.21165 From NLM Medline.
(20) Kahl, S. B.; Schaeck, J. J.; Koo, M.-S. Improved Methods for the Synthesis of Porphyrin Alcohols and Aldehydes from Protoporphyrin IX Dimethyl Ester and Their Further Modification. *The Journal of Organic Chemistry* **1997**, *62* (6), 1875–1880. DOI: 10.1021/jo961381v.
(21) Smith, K. M.; Kehres, L. A. Syntheses of methyl-devinylporphyrins related to protoporphyrin-IX. Initial studies on the mechanism of the copper (II) catalysed cyclization of 1′,8′-dimethyl-a,c-biladiene salts. *Journal of the Chemical Society, Perkin Transactions 1* **1983**, (0), 2329–2335, 10.1039/P19830002329. DOI: 10.1039/P19830002329.
(22) Hardy, B. J.; Dubiel, P.; Bungay, E. L.; Rudin, M.; Williams, C.; Arthur, C. J.; Guberman-Pfeffer, M. J.; Sofia Oliveira, A.; Curnow, P.; Anderson, J. L. R. Delineating redox cooperativity in water-soluble and membrane multiheme cytochromes through protein design. *Protein Sci* **2024**, *33* (8), e5113. DOI: 10.1002/pro.5113 From NLM Medline.
